# RNA binding protein PRRC2B mediates translation of specific mRNAs and regulates cell cycle progression

**DOI:** 10.1101/2022.12.16.520836

**Authors:** Feng Jiang, Omar M. Hedaya, EngSoon Khor, Jiangbin Wu, Matthew Auguste, Peng Yao

## Abstract

Accumulating evidence suggests that posttranscriptional control of gene expression, including RNA splicing, transport, modification, translation, and degradation, primarily relies on RNA binding proteins (RBPs). However, the functions of many RBPs remain understudied. Here, we characterized the function of a novel RBP, Proline-Rich Coiled-coil 2B (PRRC2B). Through photoactivatable ribonucleoside-enhanced crosslinking and immunoprecipitation and sequencing (PAR-CLIP-seq), we identified transcriptome-wide CU- or GA-rich PRRC2B binding sites near the translation initiation codon on a specific cohort of mRNAs in HEK293T cells. These mRNAs, including oncogenes and cell cycle regulators such as *CCND2* (cyclin D2), exhibited decreased translation upon PRRC2B knockdown as revealed by polysome-associated RNA-seq, resulting in reduced G1/S phase transition and cell proliferation. Antisense oligonucleotides blocking PRRC2B interactions with *CCND2* mRNA decreased its translation, thus inhibiting G1/S transition and cell proliferation. Mechanistically, PRRC2B interactome analysis revealed RNA-independent interactions with eukaryotic translation initiation factors 3 (eIF3) and 4G2 (eIF4G2). The interaction with translation initiation factors is essential for PRRC2B function since the eIF3/eIF4G2-interacting defective mutant, unlike wild-type PRRC2B, failed to rescue the translation deficiency or cell proliferation inhibition caused by PRRC2B knockdown. Altogether, our findings reveal that PRRC2B is essential for efficiently translating specific proteins required for cell cycle progression and cell proliferation.

**Graphic Abstract:** 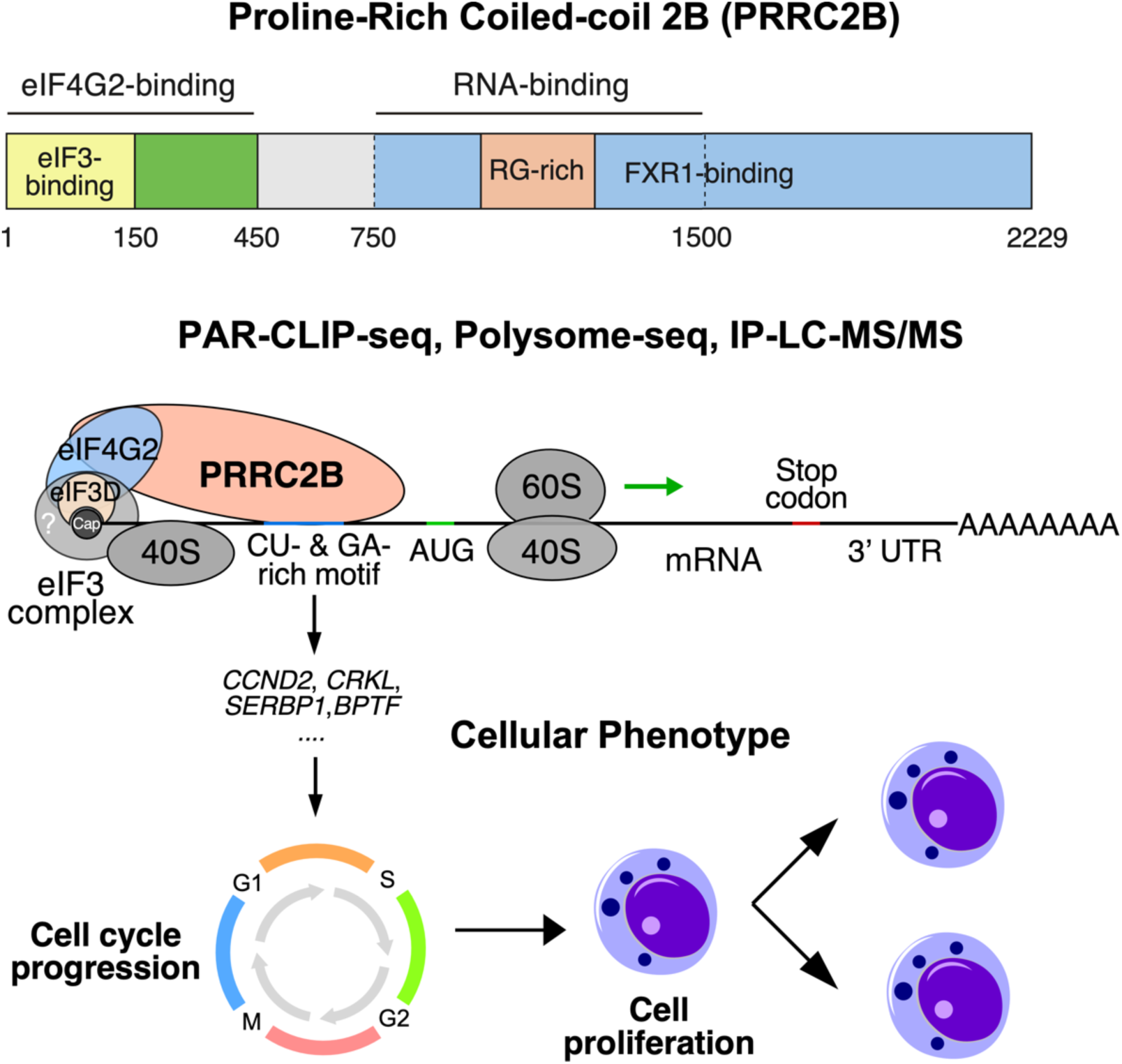

## Introduction

RNA-binding proteins (RBPs) are vital regulators affecting the fate of nearly all classes of RNA molecules (1,2). Through one or multiple RNA-binding domains (3,4), RBPs interact selectively or globally with RNAs throughout their lifespan with temporal, spatial, and functional dynamics (5). Once recruited to RNAs, RBPs participate in posttranscriptional regulation of gene expression, including RNA splicing, transport, modification, translation, and degradation (6,7). Accumulating evidence has demonstrated the essentiality of RBP-directed posttranscriptional regulations at cellular and organismal levels (7) with alterations in the abundance or functionality of RBPs linked with various human disorders (8,9). Thus, exploring the role of RBPs in posttranscriptional regulation of gene expression contributes significantly to our understanding of basic biology and human diseases.

mRNA translation is one of the posttranscriptional processes affected by RBPs extensively (10). As the most energy-consuming and precisely regulated process in cells, translation is tightly controlled through cis-acting RNA elements such as terminal oligopyrimidine (TOP) motifs and CA-rich elements (11–13) and through trans-acting factors such as mRNA binding proteins (mRBPs) (2,10,14). The latter affects all steps in global or transcript-specific translation (10). Numerous mRBPs and (m)RNA-binding domains have been identified through mass-spectrometry-based “mRNA interactome capture” methods where probes, mostly deoxythymidine oligonucleotides (oligo(dT)), were used to capture poly(A)-bearing mRNAs together with their interacting mRBPs after ultraviolet (UV) light crosslinking (15–17). These methods identify mRBPs physically bound to sequence- or structure-specific elements in mRNAs that modulate gene expression at the posttranscriptional level (6). However, despite continuous attempts at functional characterization of individual mRBPs (5), most newly identified mRBPs remain unstudied.

One of the functionally unannotated mRBPs that may regulate translation is <u>Pr</u>o-<u>r</u>ich <u>C</u>oiled-coil <u>2B</u> (PRRC2B). PRRC2B is identified as an mRNA-binding protein based on its enrichment with the oligo(dT) captured mRNAs across multiple different cell types (15,18) and its putative arginine-glycine (RG)-rich domains documented to interact with RNA (19,20). Previous studies suggested that PRRC2B is part of the eukaryotic initiation factor 4G2 (eIF4G2)-mediated translation initiation complex (21–23). This is based on its co-immunoprecipitation with eIF4G2 and eukaryotic translation initiation factor 3 (eIF3) in human breast cancer cells and mouse embryonic stem cells (21–23). However, the exact function and mRNA targets of PRRC2B in translational regulation remain unknown.

Although understudied at the mechanistic level, PRRC2B has been linked with various human cancers. PRRC2B is highly expressed in fast-proliferating tumor cells such as Wilms’ tumor, and its high expression is tightly associated with poor overall survival of patients (24,25). In addition, fusion genes of PRRC2B with MGMT, DEK, and ALK have been reported in multiple different tumors (26–30). Therefore, understanding the function of PRRC2B may shed light on translational regulation with relevance to tumor cell proliferation and human diseases.

In this study, we perform a comprehensive characterization of PRRC2B in translational regulation of gene expression in human embryonic kidney (HEK293T) cells. We demonstrate that PRRC2B and a 750 amino acid-long fragment of PRRC2B bind to GA- or CU-rich elements near the translation initiation codon of the main open reading frame on a subset of mRNAs, promoting their translation via interacting with translation initiation factors. Loss of PRRC2B binding leads to decreased translation efficiency of PRRC2B-bound mRNAs encoding oncogenes and cell cycle regulators such as cyclin D2, resulting in reduced cell cycle progression. Altogether, our study revealed the function of PRRC2B in translational regulation of specific proteins affecting cell proliferation.

## Material and Methods

### Molecular cloning

DNA fragments were obtained and amplified by PCR from Genomic DNA or cDNA extracted from HEK293T cells by Trizol (ThermoFisher) according to manufacturer’s instructions. Amplified DNA fragments were separated on 1% agarose gel, purified, digested by restriction endonucleases together with the backbone plasmid at 37°C overnight, and ligated to the cleaved backbone by T4 DNA ligase (NEB). The ligated products were transformed to *E.coli* competent cells (DH5α). Miniprep was performed on single colonies to purify the plasmids. Plasmid sequences were then confirmed by Sanger sequencing.

### Cell culture and transfections

Human HEK293T cells were propagated in Dulbecco’s modified Eagle’s medium (DMEM; Corning) supplemented with 10% fetal bovine serum. Cells were transfected with plasmids using Lipofectamine 3000 (Invitrogen) as described by the manufacturer.

### PRRC2B knockdown

shRNA cassettes were cloned into 5′-XhoI and 3′-EcoRI sites of tetracycline-inducible lentiviral pTRIPZ vector driving the expression of a TurboRFP fluorescent reporter (GE Dharmacon technology). The shRNA cassette sequences were as follows:

PRRC2B shRNA (V3THS_374329): 5′- TCATCATCACTGAACTTCA-3′
PRRC2B shRNA-328 (V3THS_374328): 5′- ATCGTTTGAGGAACCAGCC -3′
PRRC2B shRNA-626 (V2THS_159626): 5′- TTTCCCATGAACTGTGAAG-3′
Non-silencing sequence control shRNA: 5′-AATTCTCCGAACGTGTCACGT-3′

Stable single clone cell lines were generated using puromycin selection. shRNA expression was induced by 1–2 μg/ml Doxycycline and indicated as red fluorescent protein expression. Significant fluorescent signal was observed at 12 hours after addition of Dox. Induction time was counted right after addition of Dox in the cell culture media.

### Luciferase reporter construction

The 5’ UTR of *CCND2* mRNA was cloned into the pGL3-TK-5UTR-BsmBI-Luciferase reporter plasmid purchased from Addgene (https://www.addgene.org/114670/). The DNA fragment was amplified from cDNA prepared from HEK293T using primers with the extra 5’ end corresponding to the BsmbI cut sites in the plasmid: 5′-AACGTCTCCACAC-3′ for the forward primer and 5′-AACGTCTCTCTTCCAT-3′ for the reverse primer. The DNA fragment and backbone were then cleaved via the restriction enzyme BsmbI for 1 hour at 55°C and then ligated using T4 DNA ligase at room temperature, followed by selection on X-Gal coated plates. White colonies were cultured and verified using Sanger sequencing. To construct mutant 5’ UTRs, Q5 high fidelity polymerase was used to introduce the desired mutations. 0.5 μl of the PCR reaction is then incubated with a mixture of T4 DNA ligase, T4 PNK, and DpnI in T4 DNA ligase buffer for 1 hour (37°C, 15 min; 25°C,30 min; 37°C, 15 min). 5 μl of this reaction is transformed into competent cells and then plated on ampicillin agarose plates.

### PAR-CLIP

FLAG-tagged full-length and fragments of PRRC2B were cloned into 5′-KpnI and 3′-NotI sites in pcDNA3.1 plasmid for overexpression in HEK293T cells. PAR-CLIP was performed essentially following the protocol previously described (31) with some modifications. Briefly, cells were treated with 800 mM 4sU 16 hours before crosslinking by 312 nm ultraviolet exposure using Stratagene Stratalinker UV 1800. Crosslinked cells were lysed in a native lysis buffer with 50 mM HEPES, pH 7.5, 150 mM KCl, 2 mM EDTA, 1 mM NaF, 0.1% (v/v) Nonidet P-40, 1 mM DTT, 25 μl/ml protease inhibitor cocktail for mammalian tissues (Sigma-Aldrich), and 1U/μl RNase T1. Immunoprecipitation was performed using anti-DYKDDDDK magnetic agarose (ThermoFisher) followed by on-bead RNA digestion with 5U/μl RNase T1, on-bead dephosphorylation with calf intestinal alkaline phosphatase (CIP), and on-bead phosphorylation with T4 PNK and ^32^[P]-ATP. Radio-labeled RNA-protein complexes were extracted from beads by boiling in 70 μl of SDS loading buffer (Bio-Rad), separated on a 4-12% Bis-Tris gel, and imaged with phosphorimager. Bands corresponding to the protein size were cut off from the gel and subjected to proteinase K digestion and RNA purification by phenol-chloroform. The purified RNAs were run on 15% denaturing polyacrylamide gel and purified for deep sequencing. Sequencing libraries were constructed using the NEBNext® Small RNA Library Prep Set following the manufacturer’s instructions. Sequencing was performed on Illumina HiSeq2500 sequencer with 75 bp single-end sequencing.

### PAR-CLIP data analysis

We applied the CLIP Tool Kit (CTK) to perform all steps of the analysis from raw reads to binding sites and target genes as previously described (32). Briefly, raw reads were filtered based on the mean quality score with CTK built-in function and processed by cutadapt/b1 (33) to remove adapters. Processed reads with the identical sequences were labeled as PCR duplicates and removed from downstream analysis. Mapping was performed using bowtie/1.3.1 (34) with the --best –strata option against the Homo sapiens reference genome (GRCh38, hg38) downloaded from the UCSC genome browser (http://genome.ucsc.edu). One mismatch for each alignment was allowed to tolerate the T to C mutation introduced by 4sU crosslinking. Clustering and peak calling were performed on aligned reads with CTK built-in functions using the ‘valley seeking’ algorithm (32). Peak significance was first assigned based on whether the observed peak height is more than expected by chance using background models and scan statistics, then further evaluated based on the presence of T to C mutations. Significant peaks around T to C mutations were then treated as binding sites. Sequences flanking the T-to-C mutation sites (−20 to +20 nt) in the overlapped binding sites between full-length PRRC2B and P2 on mRNA exons were submitted to MEME (35) for de novo motif discovery with settings to find the longest possible motif enriched against a randomized set of sequences of the same length. Motifs with E-values < 0.05 were reported.

### RNA purification and RT-qPCR

Cells were lysed by 1000 μl of Trizol (Qiagen) and mixed with 200 μl chloroform. The mixture was spun down at 16,000 g for 10 min. RNA was precipitated from the aqueous layer by adding two volumes of isopropanol and spinning down at 16,000 g for 10 min. The pellet was washed twice with 70% ethanol, left to dry, and resuspended in nuclease-free water. DNase I was used to remove genomic DNA contamination. For quantitation of mRNA, cDNAs were prepared using iScript master mix RT Kit (Biorad) and subsequently qPCR-amplified using SYBR Primer Assay kits (Biorad). Notably, when a primer set was first used, the identity of the resulting PCR product was confirmed by cloning and Sanger sequencing. Once confirmed, melting curves were used in each subsequent PCR to verify that each primer set reproducibly and specifically generates the same PCR product. We included all the detailed information required for MIQE in supplemental information and Table S8.

### Polysome profiling and RNA extraction

1×10^9^ cells were incubated with cycloheximide (100 μg/ml; Sigma) for 10 min and harvested using a native lysis buffer with 100 mM KCl, 5 mM MgCl_2_, 10 mM HEPES, pH 7.0, 0.5% Nonidet P-40, 1 mM DTT, 100 U/ml RNasin RNase inhibitor (Promega), 2 mM vanadyl ribonucleoside complexes solution (Sigma-Aldrich (Fluka BioChemika)), 25 μl/ ml protease inhibitor cocktail for mammalian tissues (Sigma-Aldrich), cycloheximide (100 μg/ml). The lysate was spun down at 1,500 g for 5 min to pellet the nuclei. The supernatant was loaded onto a 10-50% sucrose gradient and spun in an ultracentrifuge at 150,000 g for 2 hours. The gradients were then transferred to a fractionator coupled to an ultraviolet absorbance detector that outputs an electronic trace across the gradient. Using a 60% sucrose chase solution, the gradient was pumped into the fractionator and divided equally into 9 or 10 fractions. For subsequent RNA extraction, 400 μl of each fraction was mixed with an equal amount of chloroform: phenol: isoamyl alcohol and 0.1 x of 3M sodium acetate (pH 5.2), and then spun down at 16,000 g for 10 min. The upper aqueous layer was mixed with 3 volumes of 100% ethanol and 0.1 volume of 3M sodium acetate (pH 5.2) and incubated at −20°C overnight. The solution was then spun at maximum speed for 30 min to pellet the RNA, which was washed twice with 70% ethanol and resuspended in nuclease-free water. DNase I was then added to remove genomic DNA contamination followed by another round of RNA purification. For RNA deep sequencing, RNA purified from fractions #6-10 was combined as polysome-associated RNA.

### RNA sequencing and data processing

Total and polysome-associated RNA samples were subjected to DNA removal and polyA enrichment before library construction by NGS Library Prep. Paired-end sequencing was conducted at Novogene using NovaSeq6000 S4 with a depth of 20M reads/sample. Reads were demultiplexed using bcl2fastq version 2.19.0. Quality filtering and adapter removal were performed using Trimmomatic version 0.36 (36) with the following parameters: “ILLUMINACLIP:2:30:10 LEADING:3 TRAILING:3 SLIDINGWINDOW:5:25 MINLEN:32 HEADCROP:10”. Processed/cleaned reads were then mapped to the Homo sapien reference genome (GRCh38, hg38) with Hisat version 2.1.1 (37) with the default settings. The subread-2.0.1(38) package (featureCounts) was utilized to derive gene counts using the gencode.v38 gene annotations (39) with the following parameters: “-T 10 -g gene_name -B -C -p --ignoreDup --fracOverlap 0.1”. DESeq2 version 1.38.3 (40) was used within an R-4.2.2 (41) (URL: https://www.R-project.org/) environment to normalize raw counts and identify significantly changed genes. Gene ontology analyses were performed on significantly changed genes using clusterProfiler package version 4.6 (42).

### Co-immunoprecipitation

Cells from one 80% confluent 10-cm culture dish were harvested using a native lysis buffer (300 μL) with 50 mM HEPES, pH 7.5, 150 mM KCl, 2 mM EDTA, 1 mM NaF, 0.1 % (v/v) Nonidet P-40, 1 mM DTT, and 25 μl/ ml protease inhibitor cocktail (Sigma-Aldrich) for mammalian tissues. RNasin RNase inhibitor (Promega) or RNase T1 was added for RNA-dependent and independent interactome capture. Immunoprecipitation (IP) of endogenous PRRC2B was performed using 1 μg (dilution factor 1:125) rabbit anti-human endogenous PRRC2B antibody (Invitrogen) followed by Magnetic dynabead protein A (Invitrogen) pull-down. IPs of FLAG-tagged PRRC2B and PRRC2B fragments were performed using anti-DYKDDDDK Magnetic agarose (ThermoFisher Scientific). Beads were mixed with 70 μl of SDS loading buffer (Bio-Rad) and boiled for 5 min to elute proteins for western blot and mass spectrometry analysis.

### Mass spectrometry analysis

Mass spectrometry (MS) was performed by the Mass Spectrometry Resource Lab of the University of Rochester Medical Center. Briefly, the IP elution was loaded onto a 4-12% gradient SDS-PAGE gel, run for 10 min, and stained with SimplyBlue SafeStain (Invitrogen). The stained regions were then excised, cut into 1mm cubes, de-stained, reduced, and alkylated with DTT and IAA (Sigma), dehydrated with acetonitrile, and incubated with trypsin (Promega) at 37°C overnight. Peptides were then extracted by 0.1% TFA and 50% acetonitrile, dried down in a CentriVap concentrator (Labconco), and injected onto a homemade 30 cm C18 column with 1.8 μm beads (Sepax) on an Easy nLC-1000 HPLC (ThermoFisher Scientific) connected to a Q Exactive Plus mass spectrometer (ThermoFisher Scientific) for MS. Raw data from MS experiments were mapped against SwissProt human database using the SEQUEST search engine within the Proteome Discoverer software platform, version 2.2 (ThermoFisher Scientific). The Minora node was used to determine relative protein abundance using the default settings. Percolator was used as the FDR calculator, and peptides with a q-value > 0.1 were filtered out. Biological replicates and two IPs using rabbit pre-immune IgG (negative control) were included for calculating SAINT probability. The calculation was performed by the CRAPome web tool (43) according to the user guide.

### Western blotting

Cells were lysed in RIPA buffer (Thermo Fisher Scientific), and total cell proteins were separated in a 10% denaturing polyacrylamide gel, transferred to polyvinylidene difluoride membranes (PVDF; Amersham Biosciences), probed using primary antibodies, and incubated with either a mouse or rabbit secondary antibody conjugated with horseradish peroxidase (GE Biosciences). According to the manufacturer’s suggestions, protein abundance was quantified by the Pierce BCA Protein Assay kit (Thermo, UL294765) for quantitative western blotting. If applicable, the same protein mass was loaded, and β-actin was used as an internal control.

### RNA-binding protein immunoprecipitation (RIP)

Cells from one 80% confluent 10-cm culture dish were harvested in native lysis buffer (300 μL) with 50 mM HEPES, pH 7.5, 150 mM KCl, 2 mM EDTA, 1 mM NaF, 0.1 % (v/v) Nonidet P-40, 1 mM DTT, and RNasin RNase inhibitor (Promega). Immunoprecipitation of endogenous PRRC2B was performed using 1 μg (dilution factor 1:125) rabbit anti-human PRRC2B antibody (Invitrogen) followed by Magnetic dynabead protein A (Invitrogen) pull-down. As mentioned previously, bound RNA was extracted from agarose following the same Trizol (Qiagen) extraction protocol.

### Dual-luciferase reporter assay

Control or PRRC2B knockdown HEK293T cells were grown in 96-well plates until 80% confluency. The cells were then transfected with an equal amount of experimental FLuc reporter plasmid and a control renilla luciferase (RLuc) plasmid for 18 hours. The cells were then incubated with Dual-Glo luciferase substrate (Promega) according to the manufacturer’s recommendations. The final readings of the FLuc were normalized to RLuc to obtain the relative luminescence reading. To compare between shCtrl and shPRRC2B, the relative luminescence was further normalized against reporter mRNA abundance (*FLuc*/*RLuc*).

### Cellular phenotyping

For cell proliferation, cells were seeded at 500/well in 96-well plates as biological triplicates. MTT assay was performed using MTT Cell Proliferation Kit I (Sigma-Aldrich) per the manufacturer’s recommendations every 24 hours until day 7. Cell-doubling time was calculated for cells in the log phase of the growth curve. For the cell cycle, flow cytometry was performed with Propidium iodide (PI) stained cells according to the standardized protocol (44). Biological triplicates were performed for flow cytometry. G0/G1, S, and G2/M peaks were autodetected and quantified by FlowJo, and percentages of total cells detected by flow cytometry were reported.

### Statistics

All quantitative data were presented as mean ± SD and analyzed using Excel (Microsoft Office). Statistical analyses were performed using the two-tailed Student’s *t* test with *P* < 0.05 considered significant.

## Results

### PRRC2B directly interacts with a subset of mRNAs

We first applied photoactivatable ribonucleoside-enhanced crosslinking and immunoprecipitation (PAR-CLIP) (31) to identify transcriptome-wide PRRC2B binding sites in HEK293T cells. During PAR-CLIP, RBPs were crosslinked to their bound RNAs by 4-thiouridine (4sU) and UV exposure, enriched by immunoprecipitation (IP) after RNase T1 digestion, and visualized on Bis-Tris gel by radioactive labeling of bound RNAs (**Figure 1A, S1A**). Following this method, we enriched FLAG-tagged full-length and three truncated variants of PRRC2B (P1, P2, and P3) (**Figure 1B**). While the extent of enrichment is comparable (**Figure S1B**), only full-length and 750 amino acid-long RG-rich motif containing P2 variant of PRRC2B showed significant RNA signal at the position of the protein with corresponding size (**Figure 1C**). Crosslinked RNAs (mostly 19-35 nt) were then purified from PRRC2B and P2, visualized on polyacrylamide gel (**Figure 1D**), and subjected to library construction and deep sequencing. Following an established analysis pipeline with in-house modifications, we identified a total of 1062 binding sites for full-length PRRC2B (**Table S1**) and 749 binding sites for P2 (**Table S2**) from two biological replicates that highly correlate with each other (**Figure S1C, S1D**). Among PRRC2B and P2 CLIP sites, 462 overlapped binding sites were found (**Table S1, S2**), accounting for ∼44% of the full-length binding sites and ∼62% of the P2 binding sites (**Figure 1E**). The majority of the overlapped binding sites were on mRNAs (326, ∼71%), with a small portion on long noncoding RNA (lncRNA) (62, ∼13%), miRNA (25, ∼5%), snoRNA (3, ∼1%), and other RNA species (2, < 1%) (**Figure 1F**). Similar distributions were also observed in full-length PRRC2B and P2 binding sites when analyzed separately (**Figure S1E**). Further examination of the binding sites on mRNAs revealed that 5’ untranslated region (5’ UTR) possessed the highest density (binding site per 1000 nt), followed by the coding sequence (CDS), 3’ untranslated region (3’ UTR), and intron (**Figure 1G, S1F**). The enrichment in mature mRNAs (5’ UTR, CDS, 3’ UTR) versus introns was further supported by the fact that the majority of PRRC2B are localized in the cytosol where mature mRNAs (5’ UTR, CDS, 3’ UTR) are dominant (**Figure S1G**). Interestingly, we observed a highest peak of binding sites close to the start codon region of the main open reading frames (mORFs) (**Figure 1H, S1H**). A more detailed examination reveals that majority of the peaks in 5’ UTR are at 70 nt upstream of the start codon of mORFs (**Figure 1I**). Subsequent scanning of consensus sequences in the overlapped binding sites across intact mRNAs and in the sub-regions (5’ UTR, CDS, 3’ UTR) on mRNAs by MEME (35) showed motifs that are either GA- or CU-rich (**Figure 1J, S1I**), which hardly accrue within the same binding site. Together, these results suggest that PRRC2B directly binds to a select cohort of mRNAs at the region near translation initiation sites.

**Figure 1.**
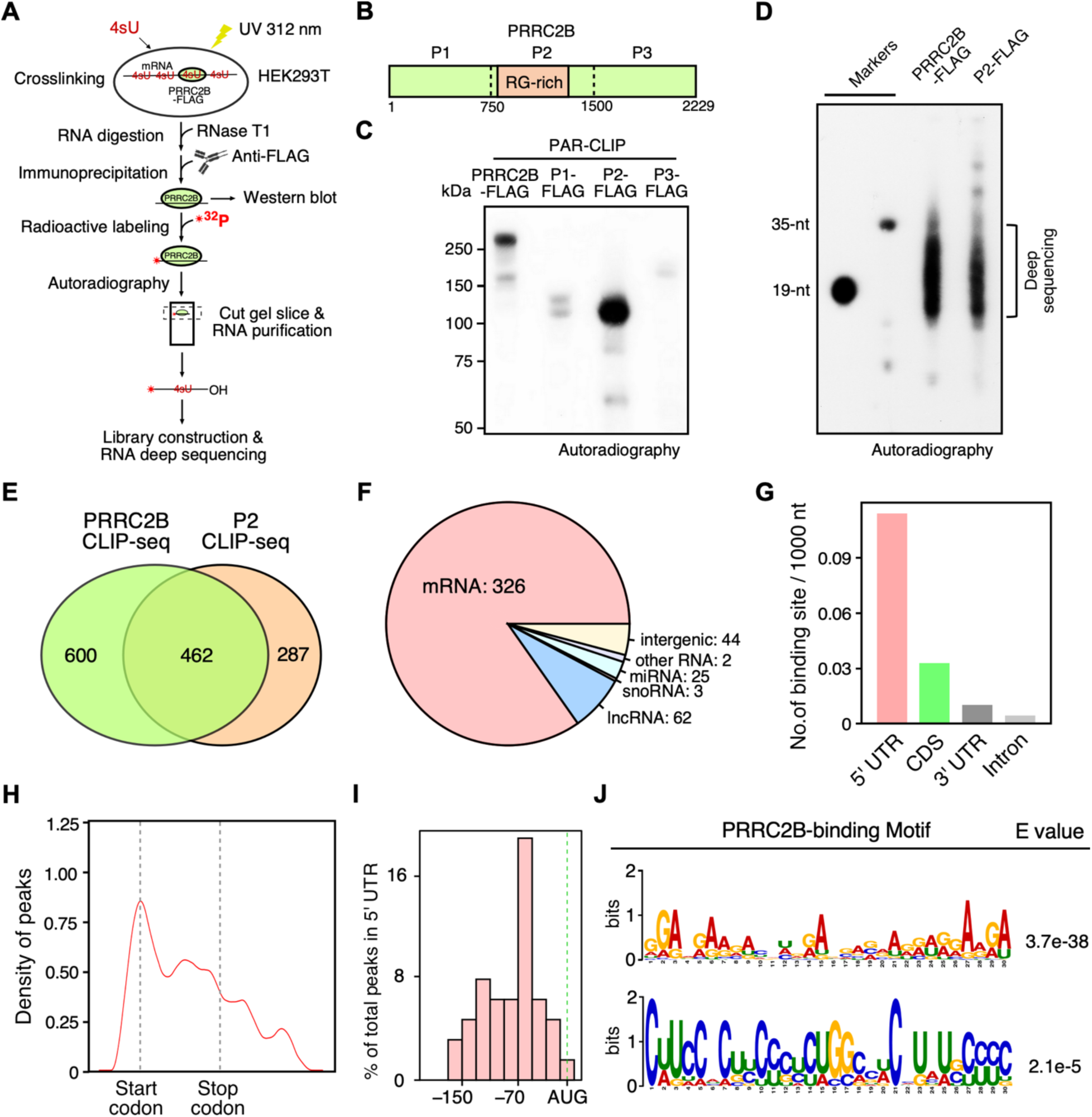
Characterization of PRRC2B-mRNA interaction through PAR-CLIP of full-length PRRC2B and its RNA-binding fragment. **(A)** A sketch of the workflow of PAR-CLIP. **(B)** Schematic showing the fragmentation of PRRC2B protein. The putative RNA-binding arginine-glycine (RG) rich sequences were covered in one fragment, P2. (**C**) An autoradiographic image showing the migration of ^32^[P]-radiolabeled crosslinked RNA-protein complexes enriched in immunoprecipitation of FLAG-tagged full-length PRRC2B and fragments (P1, P2, P3) on a 4-12% Bis-Tris gel. (**D**) Autoradiogram of ^32^[P]- radiolabeled size marker (19 and 35 nt) and RNase T1-digested RNA fragments crosslinked with full-length PRRC2B and P2 on 15% polyacrylamide gel. Results are representative of three and four independent experiments in (C, D). (**E**) A Venn diagram showing the 462 overlapped peaks (binding sites) on RNAs enriched in full-length PRRC2B and P2 immunoprecipitation. Peaks having at least 1 nt overlapping were reported as overlapped peaks. (**F**) Distribution of the 462 overlapped binding sites on different RNA species and intergenic regions. Binding sites on exons and introns were included. (**G**) A histogram showing the density of binding sites (number of binding sites per 1000 nt) on 5’ untranslated region (UTR), coding sequence (CDS), 3’ UTR, and intron regions of mRNA. Only the 326 overlapped binding sites on mRNAs were considered. (**H**) Distribution of the 326 overlapped binding sites on mature mRNAs. Only the longest transcript was included for each gene. All transcripts were scaled to the same length with a start and stop codons of the main open reading frames aligned. Kernel density estimation (KDE) was plotted as Y-axis. (**I**) Distribution of the overlapped binding sites in 5′ UTR region. The exact distance between the binding sites and start codons of the main open reading frames were plotted. (**J**) The longest significant consensus motifs that MEME can identify from the sequences flanking the T-to-C mutation sites (−20 to +20 nt) in the 326 overlapped binding sites on mRNAs. Significance was tested against randomized sequences according to the MEME user guide. E-value estimates the expected number of motifs found in a similarly sized set of random sequences (35). E-value < 0.05 was considered statistically significant.

### Loss of PRRC2B decreases translation of its bound mRNAs

The enrichment of PRRC2B binding sites near the translation initiation sites suggested its potential involvement in translational regulation. Accordingly, we examined changes in translation efficiency at the transcriptomic level upon PRRC2B knockdown by polysome profiling and RNA sequencing (**Figure 2A**) (45,46). A doxycycline (Dox)-inducible short hairpin RNA (shRNA) targeting PRRC2B (shPRRC2B) was used to knockdown PRRC2B with a non-targeting shRNA as a control (shCtrl) (**Figure 2B, 2C)**. RNA-seq was performed on polysome-associated mRNAs (associated to > 2 ribosomes, considered actively translated) and total mRNAs from cells 36 hours after Dox induction of PRRC2B knockdown to calculate translation efficiency (TE; ratio of FPKM value of polysome-associated RNA to that of total RNA) for each mRNA in PRRC2B knockdown (shPRRC2B) and control (shCtrl) cells. In the sequencing results, a total of 25537 mRNAs were detected of which 1455 mRNAs exhibited increased TE (Log_2_ (TE_shPRRC2B_/TE_shCtrl_) > 0.5; adjusted *P* value < 0.05; TE-up) upon PRRC2B knockdown while 3197 mRNAs exhibited decreased TE (Log_2_ (TE_shPRRC2B_/TE_shCtrl_) < −0.5; adjusted *P* value < 0.05; TE-down) (**Figure 2D, Table S3**). Gene Ontology (GO) analysis reveals enrichment of the TE-up genes in “pattern specification process” and “mitochondrial transport” while enrichment of TE-down genes in ‘cytosolic transport’ and ‘regulation of mRNA processing’ (**Figure S2A, S2B, Table S4**). To further elucidate the TE changes caused directly by loss of PRRC2B binding, we focused on mRNAs with the overlapped binding sites (PRRC2B-bound mRNAs) (**Table S5**). We observed significantly changed TE in 98 out of total 223 PRRC2B-bound mRNAs, among which only 18 mRNAs exhibited increased TE (Log_2_ (TE_shPRRC2B_/TE_shCtrl_) > 0.5; adjusted *P* value < 0.05; TE-up) while 80 mRNAs showed decreased TE (Log_2_ (TE_shPRRC2B_/TE_shCtrl_) < −0.5; adjusted *P* value < 0.05; TE-down) (**Figure 2E**). The TE-up PRRC2B-bound mRNAs were functionally enriched in “cytoplasmic translation” and “ribosome assembly” while TE-down PRRC2B-bound mRNAs were enriched in “stem cell population maintenance” and “maintenance of cell number” (**Figure S2C, S2D, Table S6**).

**Figure 2.**
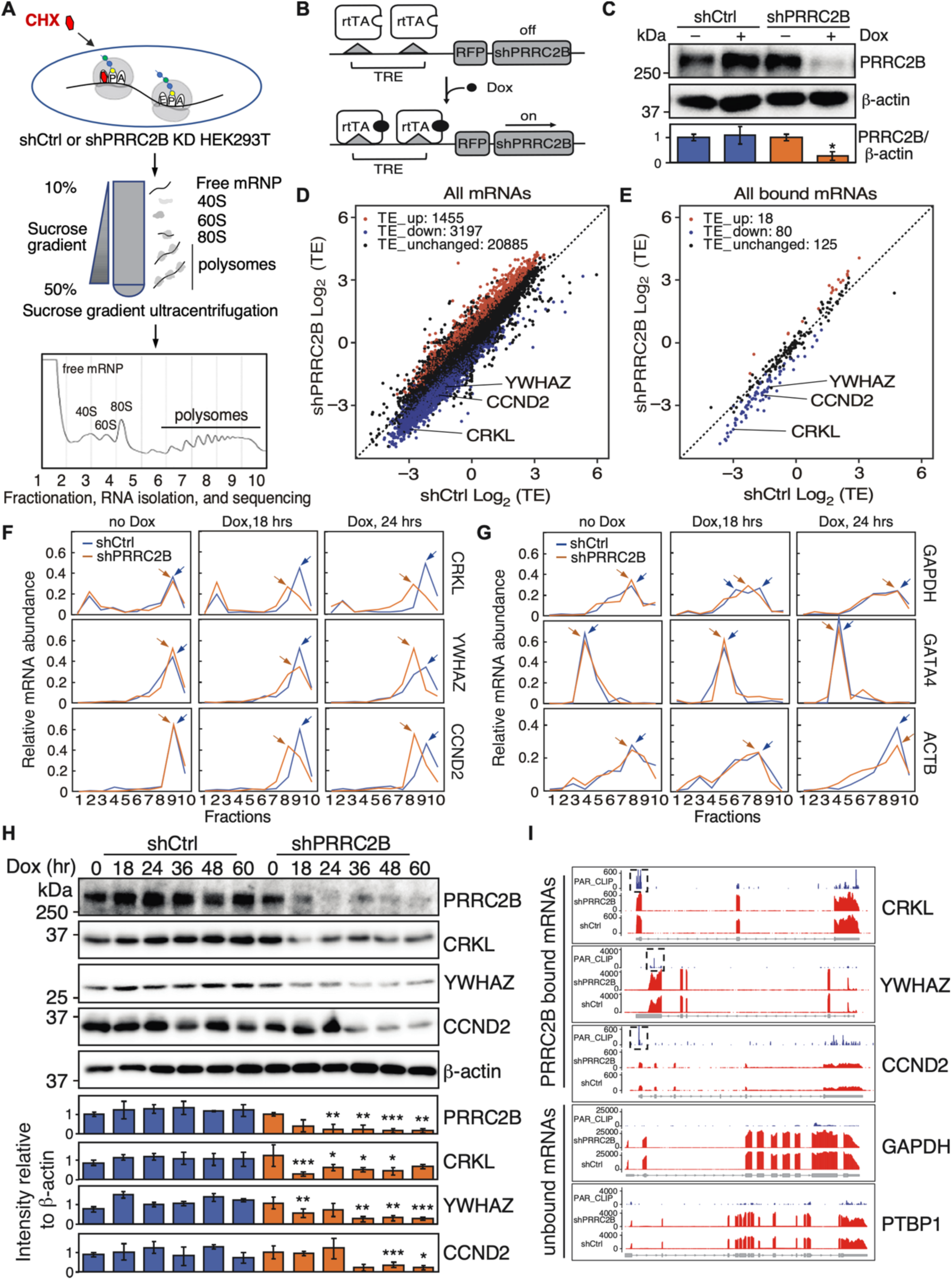
PRRC2B bound mRNAs exhibit decreased translation efficiency upon PRRC2B knockdown. (**A**) Schemes of the polysome profiling assay. Cell lysates were fractionated into 10 fractions to extract RNA for RT-qPCR. RNA sequencing was performed on the combined polysome fractions 6-10 after polysome profiling. (**B**) Schematic of the Doxycycline-inducible shRNA-mediated PRRC2B knockdown by the “Tet-on” system. TRE, tetracycline response element in Tet operator fused promoter; rtTA, reverse tetracycline-controlled trans-activator; Dox, Doxycycline; shPRRC2B, shRNA targeting the coding region of *PRRC2B* mRNA. (**C**) A western blot showed the Doxycycline-inducible knockdown of PRRC2B at 36 hours after Dox induction. shCtrl, a HEK293T cell line expressing a non-silencing control shRNA upon DOX induction; shPRRC2B, a HEK293T cell line expressing shPRRC2B upon Dox induction. (**D**) A scatterplot showing the log_2_ transformed translation efficiency (TE) values of control cells (shCtrl Log_2_ (TE)) and PRRC2B knockdown cells (shPRRC2B Log_2_ (TE)) for all mRNAs detected in polysome-seq (sequencing of pooled polysome-associated RNA) at 36 hours after Dox induction. (**E**) A scatterplot showing the log_2_ transformed translation efficiency values of control cells (shCtrl Log_2_ (TE)) and log_2_ transformed translation efficiency values of PRRC2B knockdown cells (shPRRC2B Log_2_ (TE)) for all PRRC2B bound mRNAs (mRNAs with overlapped binding sites identified in Figure 1E) at 36 hours after Dox induction. (**F, G**) mRNA abundance distribution of exemplary TE-down PRRC2B-bound mRNAs (*CRKL*, *CCND2*, *YWHAZ*), TE-unchanged PRRC2B-unbound mRNAs (*GAPDH*, *GATA4*), and TE-unchanged PRRC2B-bound mRNAs (*ACTB*) in different polysome profiling fractions (1–10) at 0, 18, 24 hours after Dox induction of PRRC2B knockdown. The distribution was measured by RT-qPCR and represented as a percentage of total RNA. Arrows indicate the peaks of mRNA abundance. (**H**) Western blot results of the protein abundance of TE-down PRRC2B-bound mRNAs (*CRKL*, *YWHAZ*, *CCND2*) at 0, 18, and 24 hours after Dox induction of PRRC2B knockdown. (**I**) Gene body coverages of PRRC2B-bound mRNAs (*CRKL*, *YWHAZ*, *CCND2*) and PRRC2B-unbound mRNAs (*GAPDH*, *PTBP1*) in PAR-CLIP of full-length PRRC2B, total RNA-seq of shCtrl cells, and total RNA-seq of shPRRC2B cells. RPKM values were plotted as Y-axis. Biological duplicates were performed for all RT-qPCR in (F, G), and representative data were shown. Western blot results are representative of >5 independent experiments in (C) and two replicates in (H). β-actin was used as an internal control in (C, H). All western blot results are presented with quantification plots and analyzed by Student’s *t* test. Significance was calculated by comparing to shCtrl (-) Dox in (C) and by comparing shPRRC2B to shCtrl at each time point in (H). * *P* < 0.05; ** *P* < 0.01; *** *P* < 0.001.

Since dysregulated TE in unbound mRNAs were detected, we suspected that secondary effects might have been triggered by the primary changes due to PRRC2B knockdown. To distinguish the primary and secondary changes, we examined the translational changes of some PRRC2B-bound and unbound mRNAs at earlier time points after Dox induction by performing reverse transcription-quantitative PCR (RT-qPCR) on RNAs extracted from polysome profiling fractions (**Figure 2A**). All tested PRRC2B-bound mRNAs were first validated to interact with the endogenous PRRC2B in cells by RNA-binding protein immunoprecipitation (RIP) and RT-qPCR (**Figure S2E**). All tested TE-down PRRC2B-bound mRNAs showed shifts towards lighter polysome or pre-polysome fractions (with smaller number of ribosomes on mRNAs) from heavier polysomes fractions (with larger number of ribosomes on mRNAs) at 18 or 24 hours after Dox induction, suggesting reduced translation efficiency (**Figure 2F, S2F**). Similar shifts were not observed in the TE-unchanged PRRC2B-bound mRNA (*ACTB*) or in PRRC2B-unbound mRNAs, including unbound TE-unchanged mRNAs (*GAPDH*, *GATA4*), unbound TE-down mRNAs (*PTBP1*, *HSPA4*), and unbound TE-up mRNA (*RPL37A*) (**Figure 2G, S2G**). This indicates that translation of PRRC2B-bound mRNAs was reduced at earlier stages (18-24 hours after Dox-induced PRRC2B knockdown) while translation of unbound mRNAs did not change significantly. Therefore, translational changes of PRRC2B-unbound mRNAs are due to secondary effects. The reduction of translation efficiency in some of the TE-down PRRC2B-bound mRNAs (*CCND2*, *CRKL*, *YWHAZ*) led to decreased protein abundance upon PRRC2B knockdown (**Figure 2H**). This observation was recapitulated using two additional shRNAs targeting PRRC2B (**Figure S2H, S2I**) and further confirmed related to translational regulation since changes (if any) at mRNA levels of *CCND2*, *CRKL*, and *YWHAZ* do not agree with protein changes (**Figure S2J**). Notably, the validated TE-down mRNAs possess PRRC2B binding sites in 5’ UTR (*CRKL*, *CCND2*), 3’ UTR (*YWHAZ*, *CTC1*, *ATAD5*), or CDS (*SERBP1*, *BPTF*, *PPP2R1A*) (**Figure 2I, S3A**), suggesting that the location of PRRC2B binding site may not affect PRRC2B function in translational control. Unlike TE-down PRRC2B-bound mRNAs, none of the tested TE-up PRRC2B-bound mRNAs showed constant shifts towards heavier polysomes (**Figure S3B**). Together, these results suggest that loss of PRRC2B primarily decreases translation of its bound mRNAs.

### PRRC2B binding to *CCND2* mRNA facilitates CCND2 protein expression

To further elucidate the role of PRRC2B binding on mRNAs, we selected cyclin D2 (*CCND2*) mRNA, which has a CU-rich PRRC2B binding site in 5’ UTR (**Figure 3A**), as an example to examine the relationship between PRRC2B binding and protein expression. Antisense oligonucleotides (ASO) base-paring with the PRRC2B binding site on *CCND2* mRNA (ASO1) were used to specifically block PRRC2B binding by forming double-stranded RNA at the PRRC2B binding site (**Figure 3A**). ASO1 significantly inhibited the binding of PRRC2B, while ASO base-paring with the region adjacent to binding sites (ASO2) (**Figure 3A**) and ASO without targets in human cells (Ctrl ASO) had minor effects (**Figure 3B**). While none of the ASOs tested caused a significant alteration in *CCND2* mRNA abundance (**Figure 3C**), ASO1 significantly decreased CCND2 protein expression in wild-type cells (**Figure 3D**). Notably, reduced CCND2 protein and mRNA expression were also observed in ASO1-treated shCtrl-expressing cells while not observed in ASO1-treated shPRRC2B-expressing cells at 36 hours after Dox induction of PRRC2B knockdown (**Figure S3C, S3D**), confirming the requirement of PRRC2B for the mechanism of action of ASO1.

**Figure 3.**
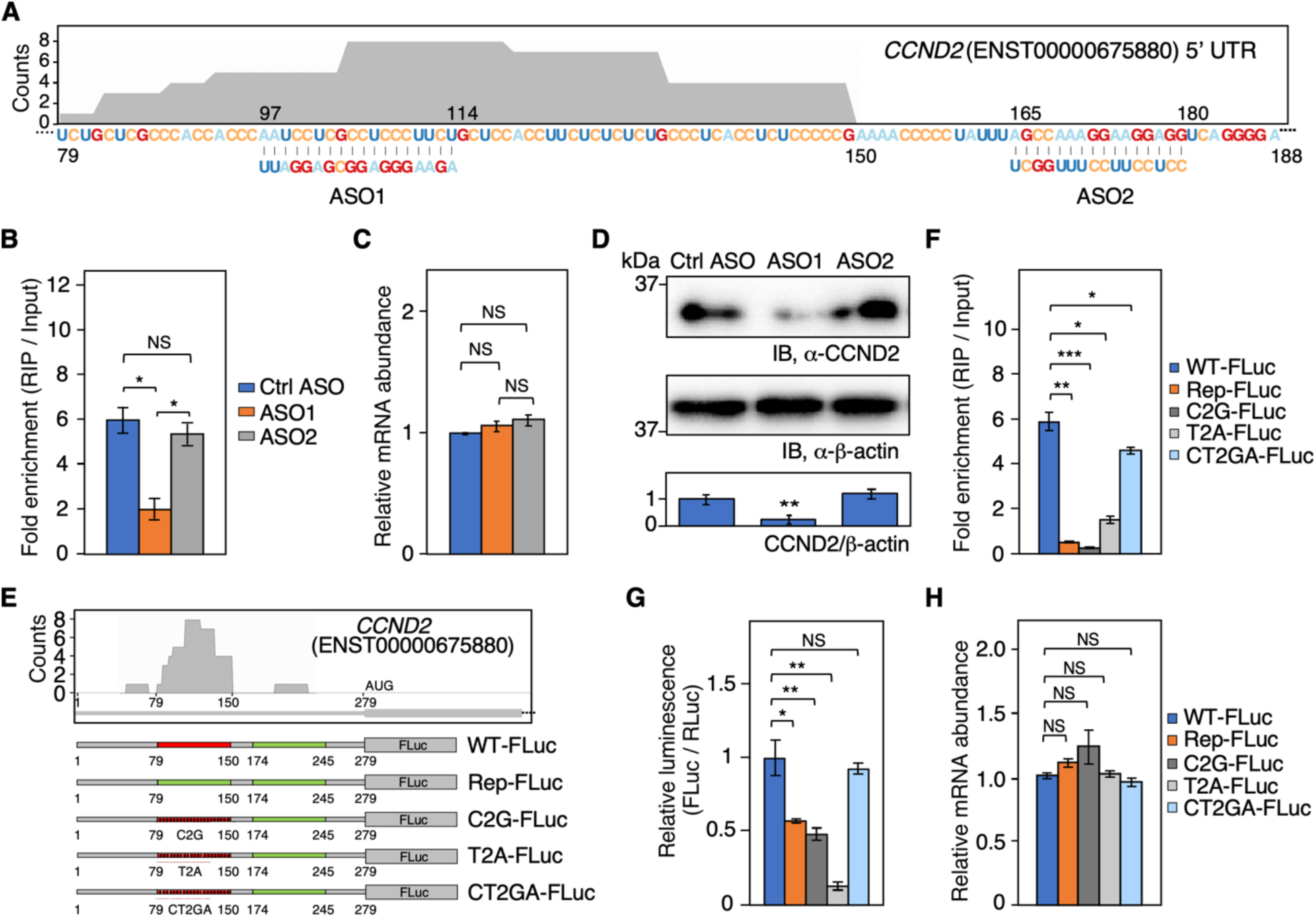
PRRC2B binding to the 5’ UTR of *CCND2* mRNA increases protein expression. (**A**) Schematic of the design of an antisense oligonucleotide (ASO) base-pairing with the PRRC2B binding site on the 5’ UTR of *CCND2* mRNA (ASO1) and the design of an ASO base-pairing with an adjacent region (ASO2). The shadow area indicates the PRRC2B binding site identified in the PAR-CLIP of PRRC2B. The height of the shadow area corresponds to the number of reads mapped to the PRRC2B binding site. (**B**) Fold enrichment (RIP versus input) of *CCND2* mRNA in RNA-binding protein immunoprecipitation (RIP) of endogenous PRRC2B in cells treated with control ASO having no targets in human cells (Ctrl ASO), ASO1, and ASO2. (**C**) Relative mRNA abundance (normalized by 18S rRNA) of *CCND2* mRNA detected by RT-qPCR in cells treated with control ASO (Ctrl ASO), ASO1, and ASO2. (**D**) Western blot analysis of the protein abundance of CCND2 in cells treated with Ctrl ASO, ASO1, and ASO2. β-actin is used as an internal control. Western blot results are representative of 3 independent experiments. (**E**) Schematic of the construction of firefly luciferase reporters with wild-type 5’ UTR of *CCND2* mRNA (WT-FLuc), 5’ UTR of *CCND2* mRNA with PRRC2B binding sites replaced by an adjacent sequence showing less PRRC2B binding (Rep-FLuc), 5’ UTR of *CCND2* mRNA with all the “C” in PRRC2B binding sites mutated to “G” (C2G-FLuc), *CCND2* 5’ UTR with all the “U” in PRRC2B binding sites mutated to “A” (T2A-FLuc), and *CCND2* 5’ UTR with all the “C & U” in PRRC2B binding sites mutated to “G & A” (CT2GA-FLuc). (**F**) Fold enrichment (RIP versus input) of reporter mRNAs in RIP of endogenous PRRC2B. (**G**) Relative luciferase activity of all reporters. (**H**) Relative mRNA abundance (normalized by 18S rRNA and *RLuc* mRNA) of reporter mRNAs detected by RT-qPCR. For all the histograms in this figure, biological triplicates were performed. Data were expressed as mean ± SD, and the Student’s *t* test was used to calculate the significance. NS, not significant; * *P* < 0.05; ** *P* < 0.01; *** *P* < 0.001. All western blot results are presented with quantification plots and analyzed by Student’s *t* test. Significance was calculated by comparing to Ctrl ASO in (D), * *P* < 0.05; ** *P* < 0.01; *** *P* < 0.001.

We next investigated the importance of the CU-rich elements in PRRC2B binding to *CCND2* mRNA by dual-luciferase reporter assays. Wild-type (WT-FLuc) and mutated 5’ UTR of *CCND2* were cloned as the 5’ UTR of the open reading frame of a firefly luciferase reporter (**Figure 3E**). These mutants include 5’ UTR of *CCND2* mRNA with (i) PRRC2B binding sites replaced by an adjacent sequence showing less PRRC2B binding (Rep-FLuc), (ii) all the “C” in PRRC2B binding sites mutated to “G” (C2G-FLuc), (iii) all the “U” in PRRC2B binding sites mutated to “A” (T2A-FLuc), and (iv) all the “C & U” in PRRC2B binding sites mutated to “G & A” (CT2GA-FLuc) (**Figure 3E, S3E**). As expected, the mRNA of WT-FLuc bound strongly with PRRC2B while the mRNAs of Rep-FLuc, C2G-FLuc, and T2A-FLuc showed significantly decreased binding affinity (**Figure 3F**). Intriguingly, the mRNA of CT2GA-FLuc only showed a moderate decrease in PRRC2B binding (**Figure 3F)**. Subsequent dual-luciferase reporter assays showed decreased luciferase activity of Rep-FLuc, C2G-FLuc, and T2A-FLuc and a modest change in luciferase activity of CT2GA-FLuc in wild-type cells, which suggests that PRRC2B binding to the reporter mRNA facilitates luciferase expression (**Figure 3G)**. These effects are further confirmed to occur at the translational level since comparable abundance of the reporter mRNAs was observed despite the sequence differences (**Figure 3H**). To test whether the observed luciferase expression differences are PRRC2B-dependent rather than just sequence-dependent, we measured the reporter expression in shCtrl cells and shPRRC2B cells at 36 hours after Dox induction of PRRC2B (**Figure S3F)**. Compared with WT-FLuc or CT2GA-FLuc, we observed that the expression of Rep-FLuc and C2G-FLuc decreased in shCtrl cells while increased in shPRRC2B cells (**Figure S3F,** orange and dark gray bar). Although the exact reason for the increased expression in shPRRC2B cells is unclear (possibly due to secondary or compensatory effect triggered after PRRC2B knockdown), our data suggests that PRRC2B plays a role in the translation regulation of *C2G-FLuc* and *T2A-FLuc* mRNAs. T2A-FLuc, on the other hand, showed little expression in both shCtrl and shPRRC2B cells (**Figure S3F, light gray bar**), suggesting that the 5’ UTR sequence changes (T2A) themselves might decrease luciferase expression independent of PRRC2B to some extent. Interestingly, we still observed significant luminescence of WT-FLuc and CT2GA-FLuc in shPRRC2B cell (**Figure S3F, blue bars**). We suspect that this might result from other mechanisms that can translate PRRC2B mRNA targets in the absence of PRRC2B and the long half-life of the luciferase protein. Together, these results suggest that PRRC2B binding to the target mRNA, *CCND2*, promotes protein expression without changing the mRNA abundance.

### PRRC2B knockdown inhibits cell proliferation

We next examined the biological consequence of PRRC2B binding on target mRNAs by focusing on cellular changes upon PRRC2B knockdown. As “maintenance of cell number” is the top functionally enriched GO term of TE-down PRRC2B-bound mRNAs (**Figure S2D, Table S6**), changes in cell proliferation were first examined. Decreased cell proliferation and increased doubling time were observed from 1 to 7 days after induction of PRRC2B knockdown (**Figure 4A, 4B**). This change could be partially explained by the observed decrease in the G1/S phase transition (**Figure 4C, 4D**). Of note, similar changes were not observed in cells without induction of PRRC2B knockdown while confirmed in cells treated with two additional shRNAs targeting PRRC2B (**Figure S4A-H**). The decreased G1/S phase transition could result from the reduced expression of cyclin D2, which has been established to facilitate the G1/S phase transition (47,48). Accordingly, decreased G1/S phase transition was observed when PRRC2B binding to *CCND2* mRNA was blocked by ASO1 (**Figure 4E, 4F**). Together, these results suggest that PRRC2B affects cell proliferation through regulating the translation of cell cycle regulators such as cyclin D2.

**Figure 4.**
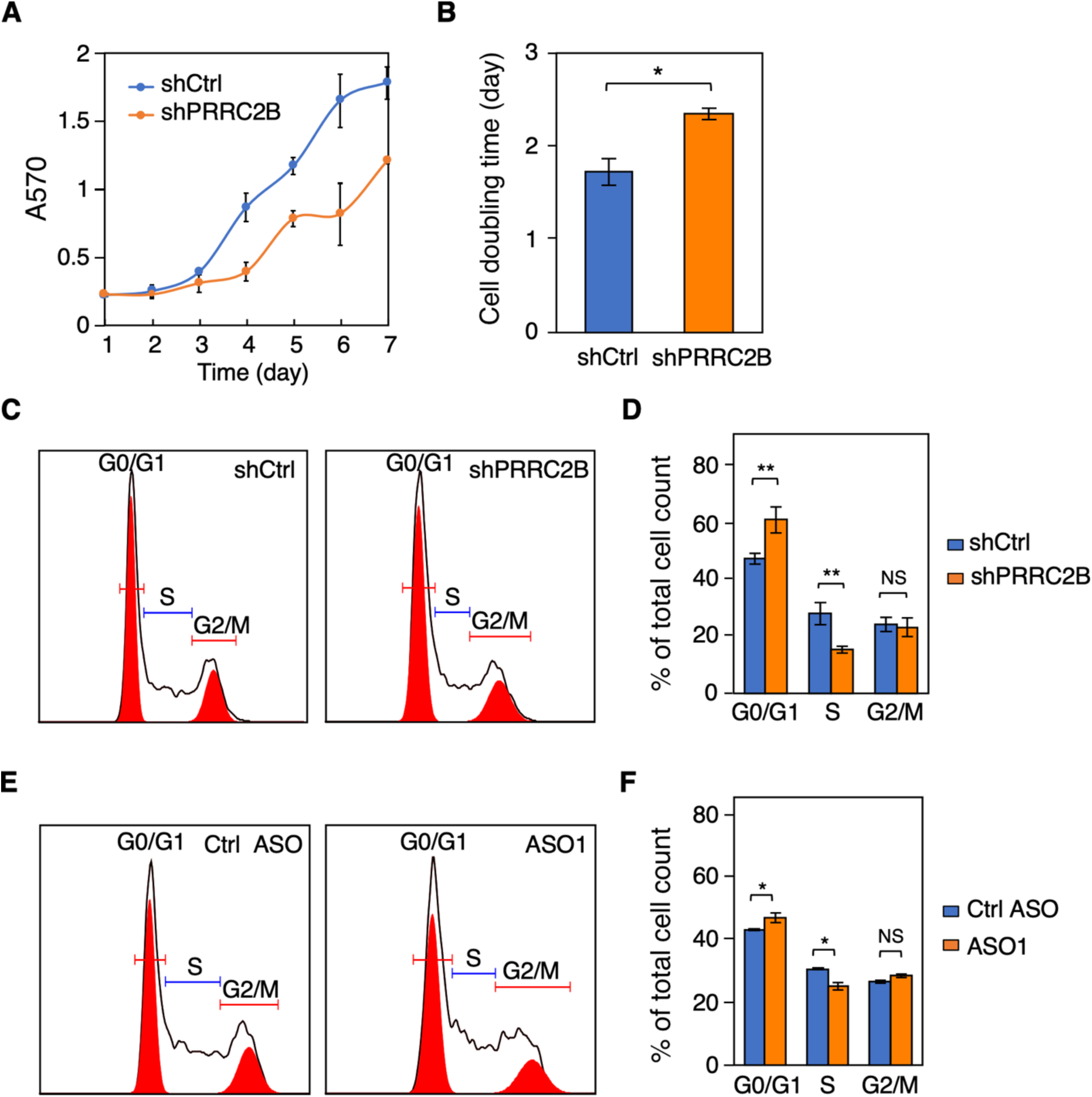
PRRC2B knockdown reduces cell proliferation and G1/S transition. (**A**) Cell proliferation curves from day 1 to day 7 of control cells (shCtrl) and PRRC2B knockdown cells (shPRRC2B) were measured by MTT assay. (**B**) Doubling time was calculated by fitting the cell proliferation data (A) to exponential functions. (**C**) Representative flow cytometry images of control cells (left, shCtrl) and PRRC2B knockdown cells (right, shPRRC2B). The experiments were performed 36 hours after Dox induction of PRRC2B knockdown. (**D**) The quantification of relative cell number across different cell cycle stages in (C). (**E**) Representative flow cytometry images of cells treated with control ASO (left) and ASO1 reducing the binding of PRRC2B to *CCND2* mRNA (right). (**F**) The quantification of relative cell number across different cell cycle stages in (E). For all flow cytometry results, biological triplicates were performed, G0/G1, S, and G2/M peaks were autodetected and quantified by FlowJo, and percentages of total cells detected by flow cytometry were reported. Biological triplicates were performed for all experiments. Data were expressed as mean ± SD in A, B, D, and F, and the Student’s *t* test was used to calculate the significance. NS, not significant; * *P* < 0.05; ** *P* < 0.01; *** *P* < 0.001.

### PRRC2B interacts with eukaryotic initiation factors eIF4G2 and eIF3

To explore mechanistically how PRRC2B regulates the translation of PRRC2B-bound mRNAs, we performed IP followed by liquid chromatography-mass spectrometry analysis to capture the interactome of endogenous PRRC2B (**Figure 5A**). Ninety-five high-confidence interactions were identified (SAINT (Significance Analysis of INTeractome) probability > 0.95) (49), of which translation initiation factors eIF4G2 and eIF3 subunits were predominant (**Figure 5B, Table S7**). The 3’ UTR-binding protein FXR1 (50) known to co-IP with eIF4G2 (23) was also detected (**Figure 5B, Table S7**). These highly confident interactions were validated by western blot following IP of either endogenous or FLAG-tagged PRRC2B (**Figure 5C, S5A**). They were further confirmed to be RNA-independent since they were retained following RNase T1 digestion of RNA to less than 35-nt long (**Figure S5B, S5C**). In contrast, no interaction of PRRC2B with eIF4G1 or cap-binding protein eIF4E was observed even though they are highly abundant and drive canonical cap-dependent translation initiation (**Figure 5B, 5C, S5A, S5C**). This suggests that PRRC2B may participate specifically in the eIF4G2-mediated translation initiation (22). This notion is further supported by the co-presence of PRRC2B, eIF4G2, eIF3, and FXR1 in the 40S and 60S fractions where translation initiation is happening (**Figure 5D**). Surprisingly, a significant amount of PRRC2B was observed in the 80S and disome fractions (**Figure 5D**). Subsequent IP experiments showed that P2 and P3 variants of PRRC2B contribute to its interaction with FXR1, while P1 is primarily responsible for interactions with eIF4G2 and eIF3 (**Figure S5D**). A more detailed IP-based mapping of truncated P1 showed that the N-terminal 1-150 amino acids of PRRC2B contain the eIF3 binding domain while the 1-450 amino acids region interacts with eIF4G2 (**Figure S5E, S5F**). Sequences in these N-terminal regions are highly conserved among PRRC2B orthologues among vertebrates (from zebrafish to humans), highlighting its functional importance (**Extended Data 1**). Together, these results suggest that PRRC2B interacts with translation initiation factors on translated mRNAs.

**Figure 5.**
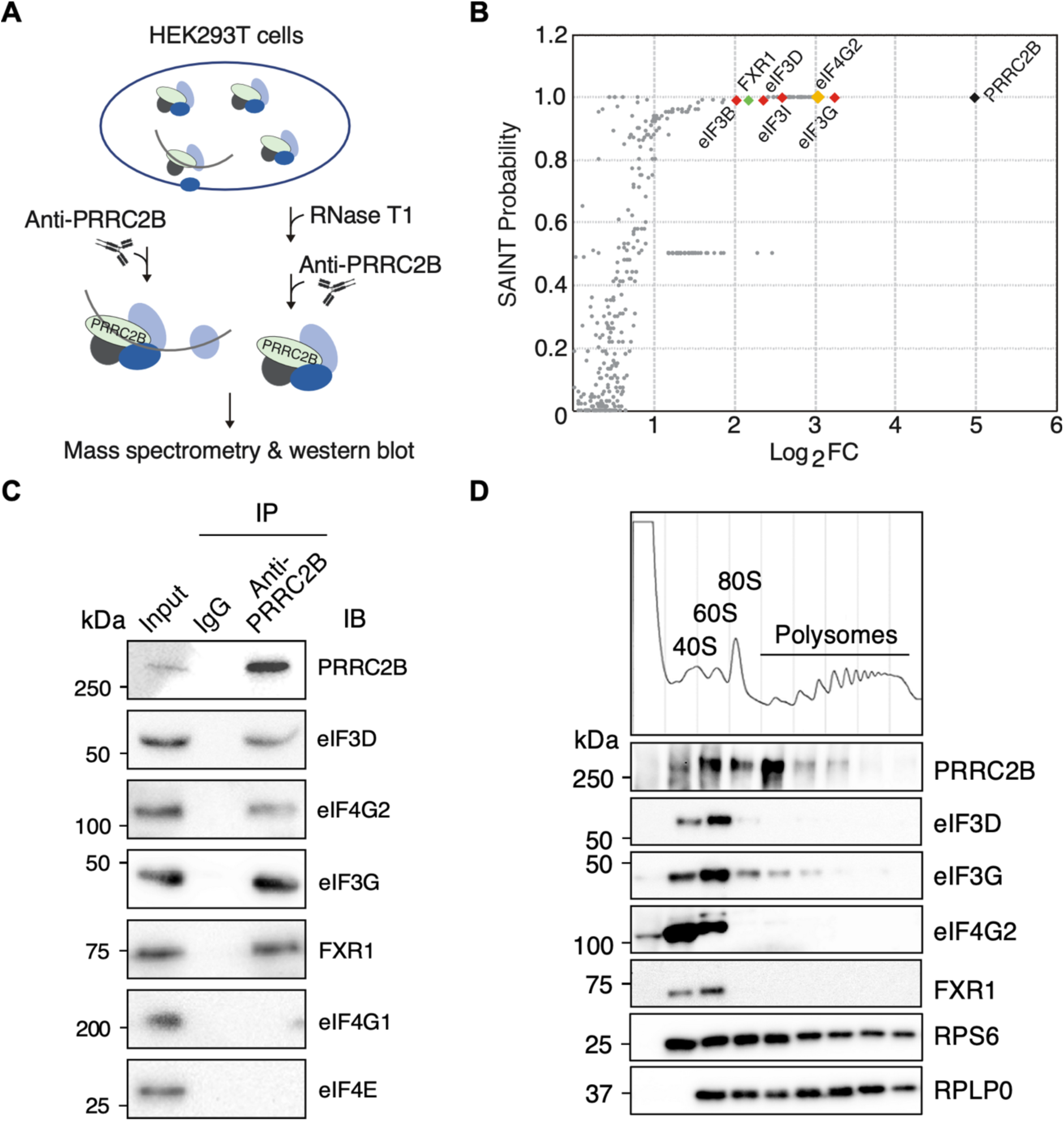
PRRC2B interacts with translation initiation factors eIF4G2 and eIF3. (**A**) Schematic of PRRC2B interactome capture workflow with or without RNase T1 digestion in HEK293T cells. (**B**) SAINT probabilities and the Log_2_ transformed fold enrichments (Log_2_FC) of all PRRC2B interacting partners identified in the PRRC2B interactome capture assay without RNase T1 treatment. Biological duplicates were performed. SAINT probability > 0.95 is considered high confidence. (**C**) Western blot analysis of the co-immunoprecipitation of endogenous PRRC2B together with translation initiation factors (eIF4G2 and eIF3) and FXR1 without RNase T1 treatment. (**D**) Western bot detection of the distribution of PRRC2B and its interacting partners in different polysome profiling fractions. Protein was purified from an equal volume of each fraction, and an equal volume of the purified protein was loaded onto a 10% SDS-PAGE gel. Western blot results represent>2 independent experiments in (C, D).

### PRRC2B functions in translational regulation by interacting with translation initiation factors eIF3 and eIF4G2

Subsequently, we examined whether PRRC2B functions through interacting with translation initiation factors eIF3 and eIF4G2. As eIF4G2 is a noncanonical translation initiation factor, we first examined its role in translating three PRRC2B-bound mRNAs (*CCND2*, *CRKL*, *YWHAZ*). We observed that all three proteins showed decreased expression upon siRNA-mediated eIF4G2 knockdown, suggesting that eIF4G2 is involved in the translation of PRRC2B-bound mRNAs (**Figure S6A**). This notion is further supported by the fact that increased expression of the three proteins was not observed in eIF4G2 knockdown cells but in wild-type cells upon overexpressing PRRC2B (**Figure 6A**). We then asked whether interacting with eIF4G2 and eIF3 is essential for the PRRC2B function. We constructed a truncated PRRC2B variant with no eIF4G2 or eIF3 binding ability (del450) (**Figure 6B, 6C**) and tested whether del450 could facilitate the translation of PRRC2B-bound mRNAs. Our rescue experiments showed that del450 failed to fully rescue the protein expression and translational changes on *CCND2*, *CRKL*, and *YWHAZ* mRNAs caused by PRRC2B knockdown, while the full length PRRC2B rescued the protein expression successfully (**Figure 6D, S6B**), suggesting del450 cannot facilitate translation of PRRC2B-bound mRNAs. Similar changes were also observed when rescuing inhibitions in cell proliferation and G1/S phase transition by wild-type PRRC2B or del450 (**Figure 6E-G, S6C**). Together, these findings demonstrate that interaction with translation initiation factors eIF4G2 and eIF3 is essential for the function of PRRC2B in translational regulation.

**Figure 6.**
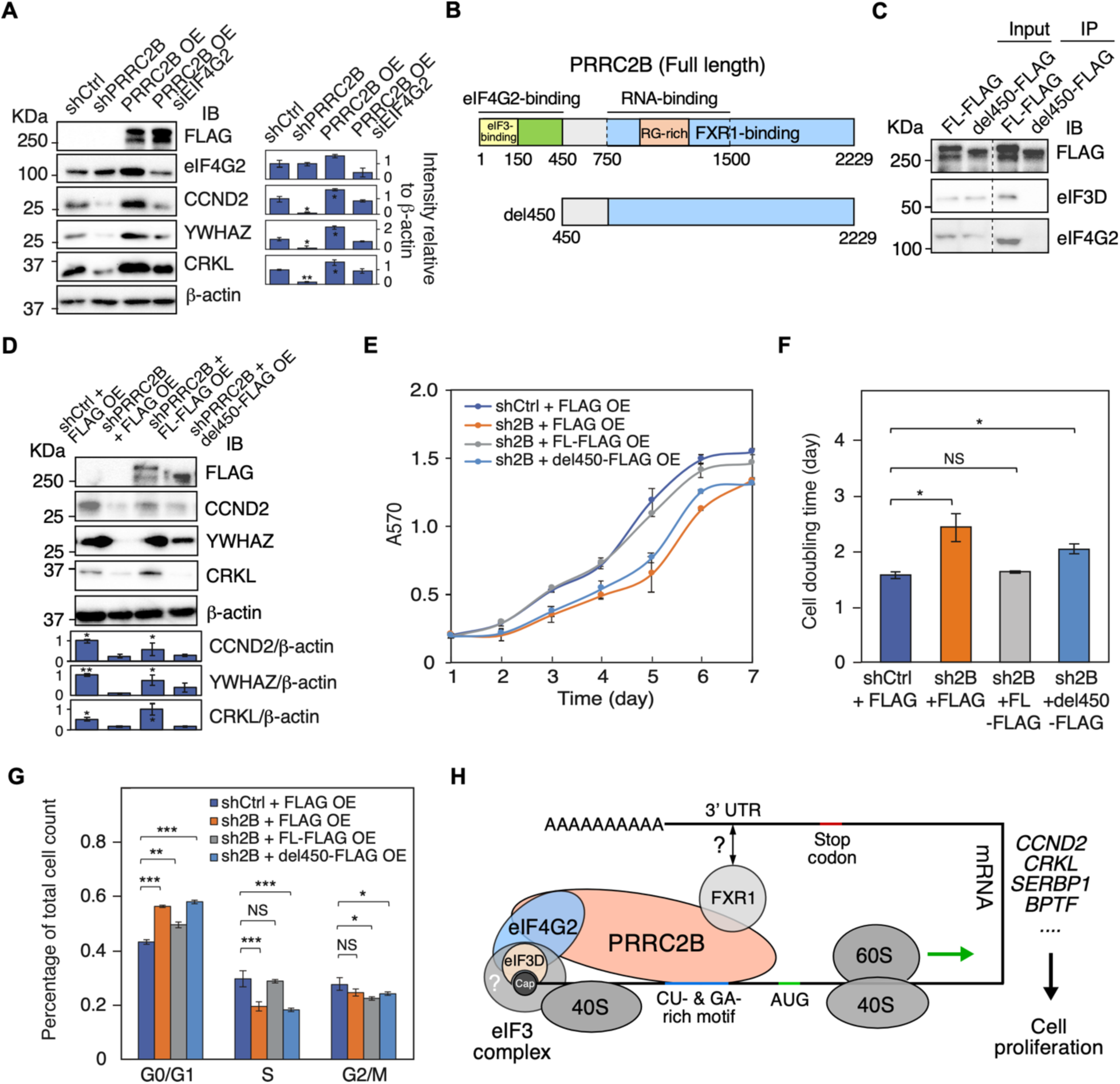
Interacting with eIF4G2/eIF3 is essential for PRRC2B-mediated translation of target mRNAs. (**A**) Western blot measurement of changes in the abundance of proteins encoded by PRRC2B-bound mRNAs (*CCND2*, *YWHAZ*, *CRKL*) in control cells (shCtrl), PRRC2B knockdown cells (shPRRC2B), PRRC2B-overexpressing cells (PRRC2B OE), and PRRC2B-overexpressing cells with eIF4G2 knockdown (PRRC2B OE siEIF4G2). (**B**) Schematic of the construction of FLAG-tagged truncated PRRC2B with eIF4G2/eIF3 binding region removed (del450). (**C**) Western blot detection of the Co-IP of FLAG-tagged full-length and truncated PRRC2B (del450) together with translation initiation factors (eIF4G2 and eIF3) upon RNase T1 treatment (5U/μl). (**D**) Western blot analysis of the protein abundance changes of CCND2, YWHAZ, and CRKL in control cells overexpressing FLAG tag (shCtrl KD + FLAG OE), PRRC2B knockdown cells overexpressing FLAG tag (shPRRC2B + FLAG OE), PRRC2B knockdown cells overexpressing FLAG-tagged PRRC2B (shPRRC2B + FL-FLAG OE), and PRRC2B knockdown cells overexpressing FLAG-tagged truncated PRRC2B (shPRRC2B + del450 OE). (**E**) Cell proliferation curves from day 1 to day 7 of cells under various conditions. (**F**) Doubling time was calculated by fitting the cell proliferation data (E) to exponential functions. (**G**) The quantification of relative cell number distribution across different cell cycle stages under various conditions. (**H**) Schematic model showing PRRC2B-bearing translation initiation complexes on mRNA. Biological triplicates were performed for all the experiments, and representative images were shown in (A, C, D). Flow cytometry results were analyzed and quantified by FlowJo. β-actin is an internal control for western blot in (A, D). Western blot results represent 3 independent experiments in (A, C, D). PRRC2B knockdown experiments were performed 36 hours after induction of gene knockdown in (A, D, G). Data were expressed as mean ± SD in (E, F, G), and the Student’s *t* test was used to calculate the significance. NS, not significant; * *P* < 0.05; ** *P* < 0.01; *** *P* < 0.001. All western blot results are presented with quantification plots and analyzed by Student’s *t* test. Significance was calculated by comparing to shCtrl in (A) and shCtrl + FLAG OE in (H). * *P* < 0.05; ** *P* < 0.01; *** *P* < 0.001.

## Discussion

In this study, we demonstrate the role of PRRC2B, a novel RNA-binding protein, in the translational regulation of PRRC2B-bound mRNAs. We discover that PRRC2B binds to CU- or GA-rich sequences near the translation initiation site, facilitating the synthesis of a group of proteins related to cell proliferation. This effect can be inhibited by antisense oligonucleotide targeting PRRC2B binding sites in PRRC2B-bound mRNAs such as *CCND2* mRNA. Mechanistically, the function of PRRC2B primarily depends on its interaction with eIF4G2 and eIF3, suggesting a positive regulatory role of PRRC2B in eIF4G2-mediated translation. Altogether, this study reveals a novel translation regulatory pathway that maintains efficient cell proliferation and can be targeted to diminish cell cycle progression.

Using PAR-CLIP (31), we demonstrated the binding of PRRC2B to mature mRNAs. Unlike many promiscuous RNA-binding proteins, PRRC2B binds to both CU- and GA-rich RNA elements. Although CU- and GA-rich RNA sequences tend to be complementary, we assume that PRRC2B binds to single-stranded RNA since these elements seldom appear adjacent to each other, and therefore are unlikely to base pair to form double-stranded RNA. Our mutagenesis assays demonstrated that mutating a CU-rich binding site to GA-rich has a minor effect on PRRC2B binding and function. We conclude that PRRC2B binds to both motifs with a similar affinity. Although CU- and GA-rich motifs are found throughout mature mRNAs (in 5’ UTR, CDS, and 3’ UTR), PRRC2B predominately binds the motif sequence near the translation initiation sites where it interacts with translation initiation factors eIF4G2 and eIF3. Despite the fact that eIF4G2 has been accounted for transcript selectivity (23,51), interacting with it hardly affects PRRC2B binding since the PAR-CLIP for P2, the 751-1500 amino acid-containing fragment of PRRC2B not interacting with eIF4G2 or eIF3, exhibited similar binding profile and target sequences as the full-length PRRC2B. Based on this, we conclude that the P2 region of PRRC2B dictates at least some degree of transcript selectivity in an eIF4G2-and eIF3-independent manner.

eIF4G2 has been evident to drive noncanonical eIF4G1- and eIF4E-independent translation initiation (22,23,52) together with eIF3D, which replaces eIF4E in cap-binding (52), and FXR1, which binds to mRNA 3’ UTR and facilitates mRNA circularization (50). We conclude from the PRRC2B interactome (**Figure 5, S5**) that PRRC2B selectively participates in eIF4G2-mediated translation initiation but not the canonical eIF4G1-mediated translation initiation. This discovery echoes previous studies on eIF4G2 (22,23) and is essential for understanding the PRRC2B function (**Figure 6**). We speculate that PRRC2B may act as a scaffold recruiting other factors since it interacts with mRNA, eIF4G2, eIF3, and FXR1 through distinct regions with little dependency on each other (**Figure 1, S5**). Besides, considering that PRRC2B dictates transcript selectivity, it may navigate eIF4G2 to specific mRNAs. However, given the mRNA abundance of *eIF4G2* is roughly three times of *PRRC2B* in HEK293T cells (FPKM value detected by our RNA-seq), PRRC2B likely only participates in a fraction of eIF4G2-mediated translation. In addition, PRRC2B does not completely co-sediment with translation initiation complexes in the 40S to 80S fractions (**Figure 5D**), suggesting that it may have other functions beyond translation initiation (e.g., initiation-to-elongation transition). To establish the role of PRRC2B in eIF4G2-mediated translation initiation, future studies involving *in vitro* translation are required. However, obtaining recombinant PRRC2B will be significantly challenging given that full-length PRRC2B is 243 KDa and highly disordered, as predicted by AlphaFold (53).

Although the mechanism is not fully defined, PRRC2B facilitates the eIF4G2-mediated translation of its bound mRNAs (**Figure 2, 3**). These mRNAs include multiple oncogenes and cell cycle regulators such as *CCND2* (47,48) and *CRKL* (54–56), agreeing with the link between PRRC2B and various types of human cancer (24–30). Our results suggest that PRRC2B promotes cell proliferation by facilitating the translation of *CCND2* mRNA, which encodes the D-type cyclin facilitating G1/S transition in cancer and stem cells (47,57). Blocking PRRC2B binding to *CCND2* mRNA by ASOs significantly decreases the CCND2 expression in cells expressing PRRC2B (**Figure 3**), thereby inhibiting cell proliferation. This discovery inspired us to inhibit the expression of oncogene and cell cycle proteins by ASOs exclusively in cells highly expressing PRRC2B. Because PRRC2B is highly expressed in cancer cells such as large B cell lymphoma (DLBC), thymus cancer (THYM) (25), and Wilms’ tumor (24), these ASOs can be applied for potential anti-cancer treatment, which deserves future investigation in biological or disease-relevant models. In addition, several other validated PRRC2B target mRNAs (**Figure S2E, F**) encode oncogenic or pro-proliferation proteins that may contribute to PRRC2B-mediated regulation of cell cycle progression, including SERBP1 (58), CTC1 (59), ATAD5 (60), and BPTF (61). Moreover, multiple Torin 1-sensitive mTORC1 (mammalian target of rapamycin complex 1) pathway downstream mRNAs (62) were identified as PRRC2B-bound target transcripts by PAR-CLIP-seq and polysome-seq, such as *EEF2*, *HSP90AB1*, and *HSPA8* (**Figure 2E, Table S5**), indicating potential crosstalk between PRRC2B-eIF4G2 and mTORC1 pathways, which may need further studies.

While PRRC2B is evident to facilitate the translation of its bound mRNAs, we still found targets to have unchanged or increased translation efficiency upon PRRC2B knockdown. The unaltered TE could indicate that the translation of PRRC2B-bound mRNAs may not entirely depend on PRRC2B-mediated translation, as there could be other mechanisms sufficient to carry out the translation when PRRC2B is absent. The increased TE is probably due to secondary or compensatory effects since these mRNAs, similar to the TE-changed unbound mRNAs, exhibited no translational changes at earlier time points after induction of PRRC2B knockdown (**Figure S2G**). These secondary or compensatory effects could result from the primary changes in TE of PRRC2B-bound mRNAs or the altered cell cycle progression upon PRRC2B knockdown (**Figure 4**) since translation varies dramatically in different phases of the cell cycle (63).

The PRRC2 proteins have been repeatedly linked with mRNA binding and posttranscriptional regulation (21-23,64-67). A recent study suggests that PRRC2 proteins facilitate leaky scanning of upstream open reading frames (uORF) (67). However, our results showed both mRNAs with and without uORFs are bound and regulated by PRRC2B, suggesting that the effect of PRRC2B is not limited to uORFs but determined by the presence of GA- and CU-rich binding sites. Besides, previous studies showed PRRC2A interacts with N6-methyladenosine (m^6^A) around the stop codon to stabilize mRNA (64), while PRRC2C facilities the efficient formation of stress granules (65,66). Although PRRC2B shares sequence similarities with PRRC2A and PRRC2C (**Extended Data 2**), the reported function of PRRC2A and PRRC2C were not recapitulated in PRRC2B (**Figure 1H, S1H, 2I, S2H, data not shown**). On the other hand, PRRC2B loss-of-function mutations have been linked with human congenital heart defects (68) and a variety of mouse developmental defects, including cardiovascular developmental defects (69). Therefore, the functional diversity and redundancy of the three members of the PRRC2 protein family warrant future elucidation.

## Data availability

Data supporting the findings of this manuscript are available from the corresponding author upon request. The sequencing data raw files and processed files were deposited in the GEO database and are accessible under accession numbers GSE220057, GSE220058, and GSE220059.

## Acknowledgments

We are grateful for the consulting contributions from Elizabeth Grayhack, Joshua Munger, Yitao Yu, and Chen Yan. We appreciate Jared Hollinger and Huan Liu for their technical support and Emily Bonanno for her polishing the manuscript. This work was supported in part by the National Institutes of Health (R01 HL132899 and R01 HL147954 to P.Y.), NIH T32 Fellowship (T32 GM068411 to O.H.), American Heart Association Career Development Award (848985 to J.W.), and start-up funds from Aab Cardiovascular Research Institute of University of Rochester Medical Center (to P.Y.).

## Author contributions

PY obtained the funding and launched the study. PY, JW, and FJ conceived the ideas. PY and FJ designed the experiments, analyzed the data, and wrote the manuscript. FJ carried out the experimental work and bioinformatic analysis. OH, ESK, and MA provided conceptual feedback or technical assistance. All the authors discussed the results and had the opportunity to comment on the manuscript.

## Conflict of interest

None of the authors have any financial conflict of interest related to the research described in this manuscript.

## Supplementary Information for

**This document file includes the following:**

Figures S1 to S6

Tables S1 to S8

Extended Data 1 and 2

SI Materials and Methods

MIQE for RT-qPCR

### Supplemental figures

**Figure S1.**
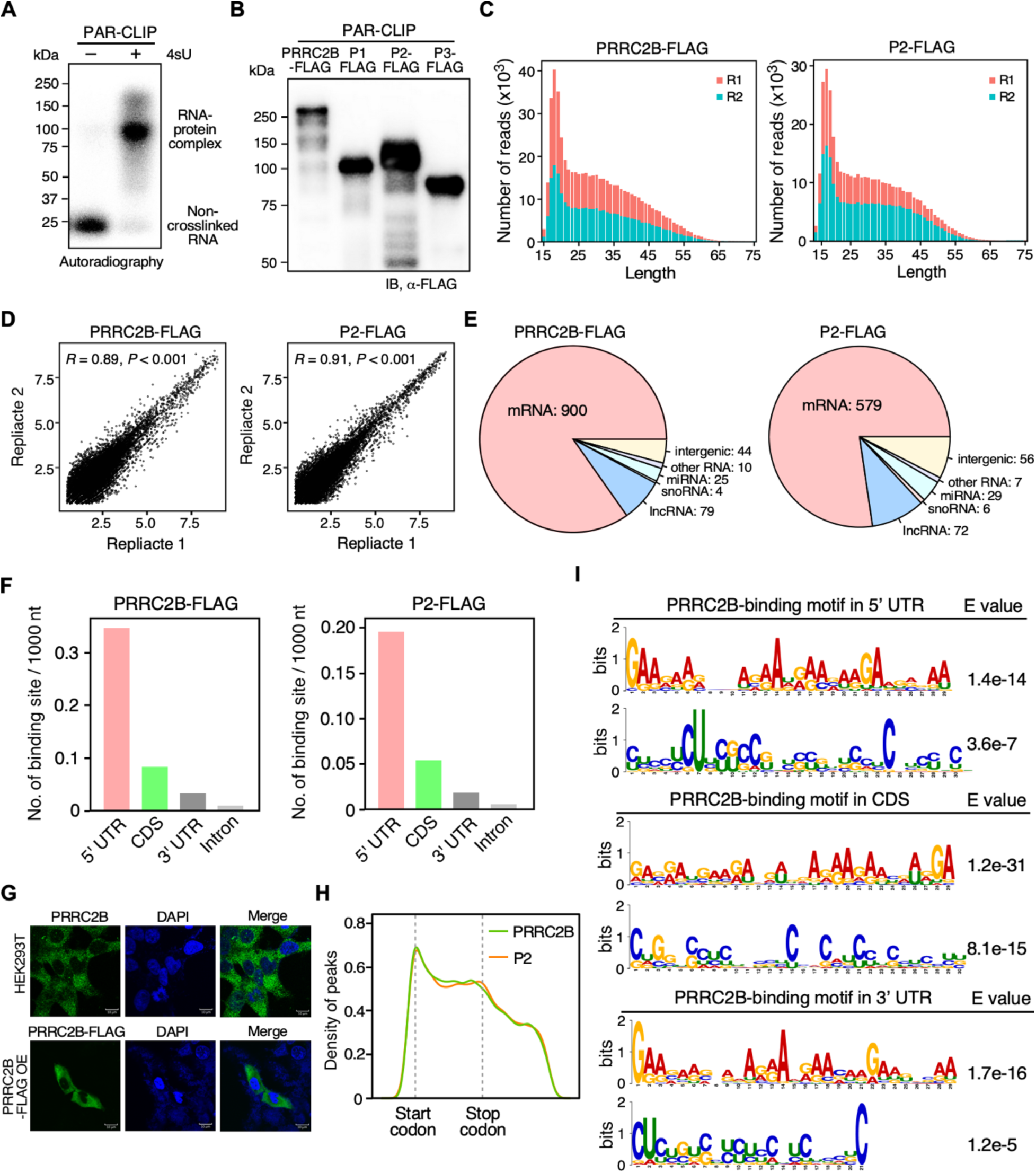
PAR-CLIP analysis of PRRC2B-RNA interaction. (**A**) An autoradiographic image showing the migration of ^32^[P]-radiolabeled crosslinked RNA-protein complexes (with 4-thiouridine, 4sU) and non-crosslinked RNAs (without 4sU) enriched in the immunoprecipitation step of PAR-CLIP on a 4-12% Bis-Tris gel. (**B**) Western blot analysis shows the enrichment of comparable protein amounts of FLAG-tagged full-length PRRC2B and fragments (P1, P2, P3) by anti-FLAG antibody-conjugated magnetic beads in PAR-CLIP. 2% of the total volume of enriched proteins was loaded on a 10% SDS-PAGE gel. (**C**) Distribution of the length of sequenced reads after trimming and quality filtering of the two biological replicates in full-length PRRC2B (left) and P2 PAR-CLIP (right). (**D**) Strong correlations (Pearson’s correlation) between the two biological replicates in full-length PRRC2B (left) and P2 PAR-CLIP (right), as indicated by scatterplots. The coordinates of each dot represent the FPKM (fragments per kilobase of exon per million mapped fragments) values of each gene in the two biological replicates. (**E**) Distribution of the binding sites identified in full-length PRRC2B (left) and P2 PAR-CLIP (right) on different RNA species and intergenic regions. Binding sites on exons and introns were included. (**F**) Histograms showing the density of binding sites (number of binding sites per 1000 nt) on 5’ UTR, CDS, 3’ UTR, and intron regions of mRNA in full-length PRRC2B (left) and P2 PAR-CLIP (right). Only the binding sites on mRNAs were considered. (**G**) Subcellular localization of endogenous and FLAG-tagged PRRC2B in HEK293T cells detected by immunofluorescent staining. (**H**) Distribution of binding sites identified in full-length PRRC2B (left) and P2 PAR-CLIP (right) on mature mRNAs. Only the longest transcript was included for each gene. All transcripts were scaled to the same length with a start and stop codons of the main open reading frames aligned. Kernel density estimation (KDE) was plotted as Y-axis. (**I**) The longest significant consensus motifs that MEME can identify from the sequences flanking the T-to-C mutation sites (−20 to +20 nt) in the overlapped binding sites on 5’ UTR, CDS, and 3’ UTR. Significance was tested against randomized sequences according to the MEME user guide. E-value estimates the expected number of motifs found in a similarly sized set of random sequences (32). E-value < 0.05 was considered statistically significant.

**Figure S2.**
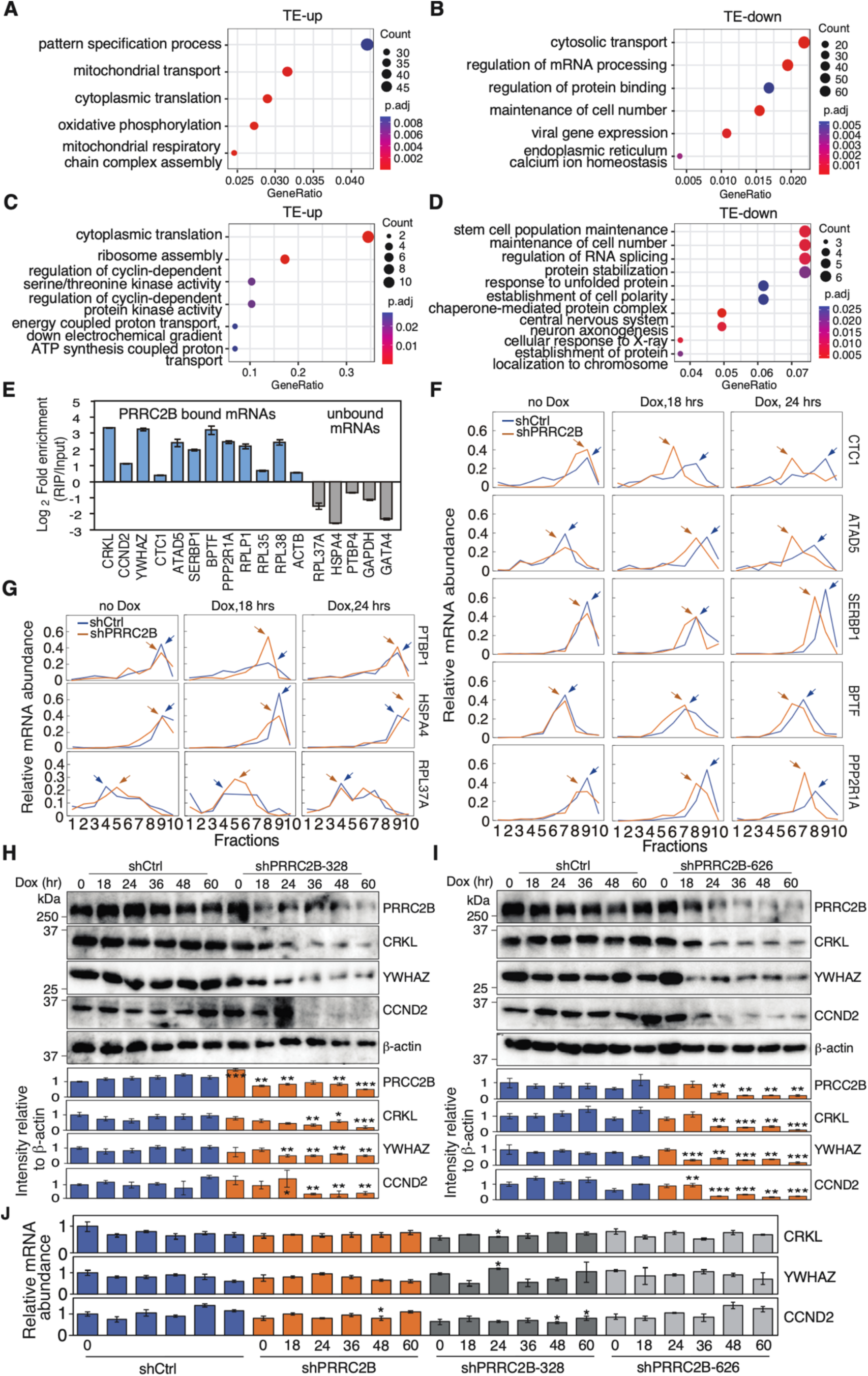
PRRC2B-bound mRNAs exhibit decreased translation efficiencies upon PRRC2B knockdown. (**A-D**) Top enriched gene ontology (GO) biological processes in all TE-up mRNAs, TE-down mRNAs, TE-up PRRC2B-bound mRNAs, and TE-down PRRC2B-bound mRNAs. GO terms were filtered by adjusted *P* < 0.05 and simplified by the R package ‘ClusterProfiler’ to remove redundant terms. (**E**) RNA-binding protein immunoprecipitation (RIP) of endogenous PRRC2B followed by RT-qPCR for PRRC2B-bound and unbound mRNAs. Input mRNA was used as a normalizer. Results are presented as mean ± SD. (**F, G**) mRNA abundance distribution (detected by RT-qPCR and represented as a percentage of total RNA) of exemplary TE-down PRRC2B-bound mRNAs (*CTC1*, *ATAD5*, *SERBP1*, *BPTF*, *PPP2R1A*), TE-down unbound mRNAs (*PTBP1*, *HSPA4*), and TE-up unbound mRNAs (*RPL37A*) in different polysome profiling fractions at 0, 18, 24 hours after Dox induction of PRRC2B knockdown. Arrows indicate the peak of mRNA abundance. Biological duplicates were performed for RT-qPCR experiments included in (F, G), and representative data were shown. (**H, I**) Western blot results of the protein abundance of TE-down PRRC2B-bound mRNAs (*CRKL*, *YWHAZ*, *CCND2*) at 0, 18, and 24 hours after Dox induction of PRRC2B knockdown by two additional shRNAs. (**J**) RT-qPCR measurement of *CRKL*, *YWHAZ*, *CCND2* mRNA in control and PRRC2B knockdown cells at different time points after Dox induction. All western blot results are presented with quantification plots and analyzed by the Student’s *t* test. Significance was calculated by comparing shPRRC2B to shCtrl at each time point. * *P* < 0.05; ** *P* < 0.01; *** *P* < 0.001.

**Figure S3.**
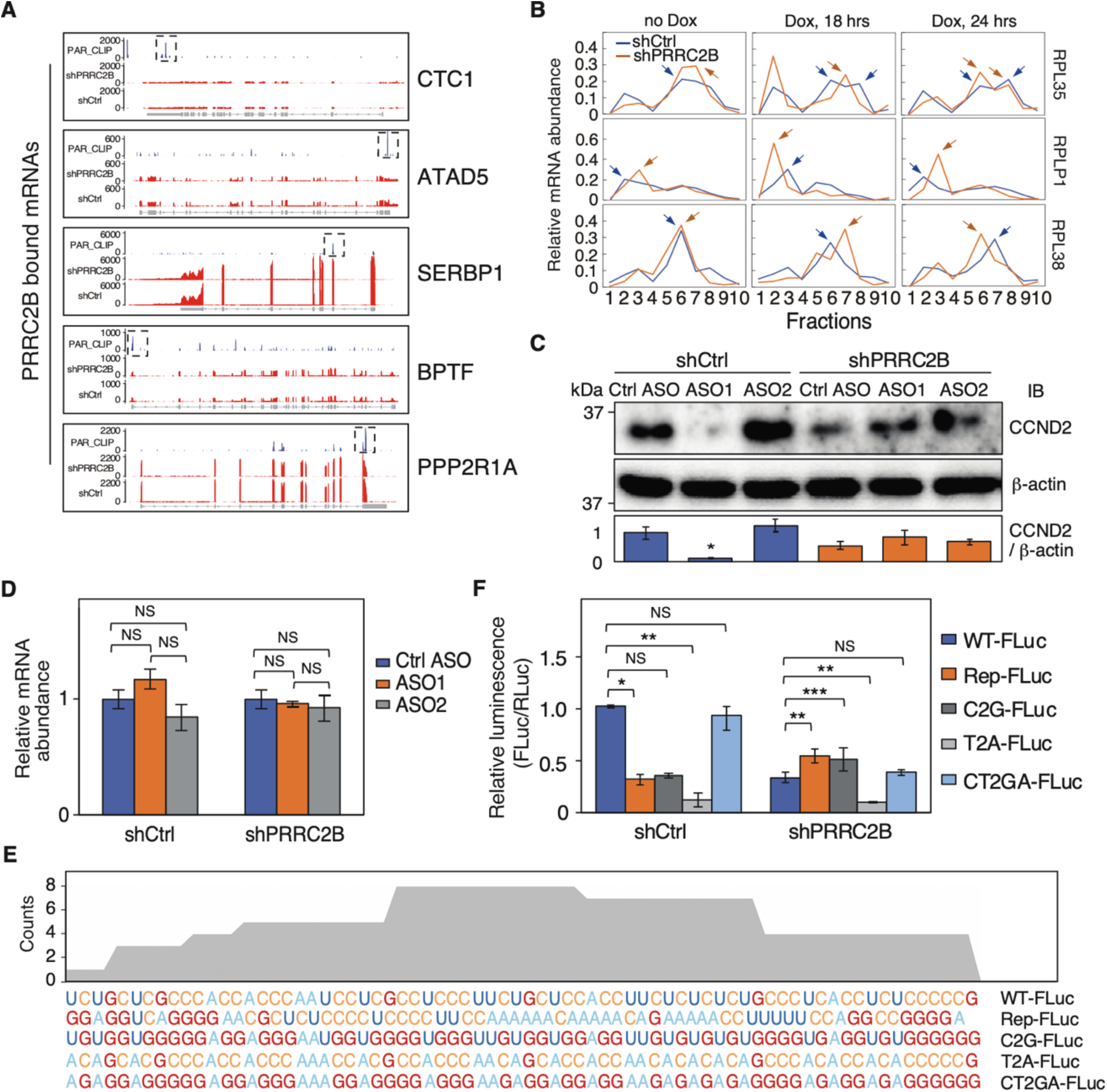
PRRC2B knockdown cells are insensitive to ASOs and express the mutant luciferase reporters differently from wild-type cells. (**A**) Gene body coverages (exhibited as FPKM values) of exemplary TE-down PRRC2B-bound mRNAs (*CTC1*, *ATAD5*, *SERBP1*, *BPTF*, *PPP2R1A*) in PAR-CLIP of full-length PRRC2B, total RNA-seq of control cells, and total RNA-seq of PRRC2B knockdown cells. RPKM values are plotted as Y-axis. (**B**) mRNA abundance distribution (detected by RT-qPCR and represented as a percentage of total RNA) of exemplary TE-up PRRC2B-bound mRNAs in different polysome profiling fractions at 0, 18, and 24 hours after Dox induction of PRRC2B knockdown. Arrows indicate the peak of mRNA abundance. Biological duplicates were performed, and representative data were shown. (**C**) Western blot analysis of the protein abundance of CCND2 in shCtrl and shPRRC2B cells treated with Ctrl ASO, ASO1, and ASO2. (**D**) RT-qPCR measurement of *CCND2* mRNA in shCtrl and shPRRC2B cells treated with Ctrl ASO, ASO1, and ASO2. (**E**) A snapshot showing the sequences of the PRRC2B binding region on WT-FLuc and corresponding mutated sequences in Del-FLuc, C2G-FLuc, T2A-FLuc, and CT2GA-FLuc. (**F**) Relative luminescence (FLuc/RLuc) of wild-type and four mutant reporters in shCtrl and shPRRC2B cells. To compare the luciferase translation between shCtrl and shPRRC2B, relative luminescence (FLuc/RLuc) was normalized against mRNA abundance (*FLuc*/*RLuc*) and presented as Y-axis values. For all assays involving PRRC2B knockdown, experiments were performed 36 hours after DOX induction. Western blots were representative of >2 biological replicates. Data were expressed as mean ± SD in (C, D, F), and the Student’s *t* test was used to calculate the significance. NS, not significant; * *P* < 0.05; ** *P* < 0.01. All western blot results are presented with quantification plots and analyzed by the Student’s *t* test. Significance was calculated separately in shCtrl and shPRRC2B cells by comparing ASO1 and ASO2 to Ctrl ASO in (C, D). * *P* < 0.05; ** *P* < 0.01; *** *P* < 0.001.

**Figure S4.**
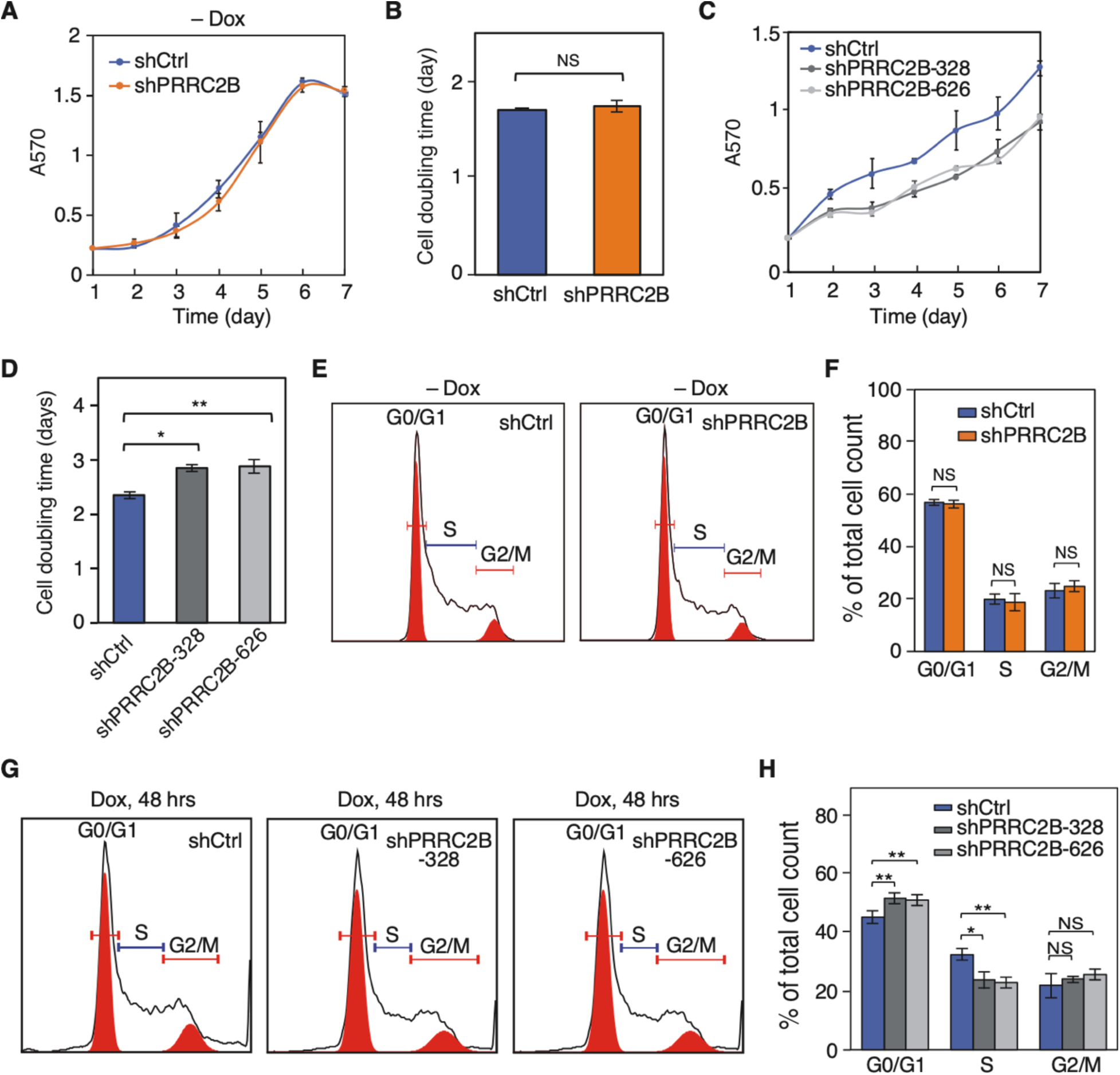
Cell proliferation or G1/S transition does not change without induction of PRRC2B knockdown. (**A**) Cell proliferation curves from day 1 to day 7 of control cells (shCtrl) and PRRC2B knockdown cells (shPRRC2B) without induction of PRRC2B knockdown (no DOX treatment) measured by MTT assay. (**B**) Doubling time was calculated by fitting the cell proliferation data (A) to exponential functions. (**C**) Cell proliferation curves from day 1 to day 7 of control cells (shCtrl) and PRRC2B knockdown cells (shPRRC2B-328, shPRRC2B-626) with induction of PRRC2B knockdown (DOX treatment) measured by MTT assay. (**D**) Doubling time was calculated by fitting the cell proliferation data (C) to exponential functions. (**E**) Representative flow cytometry images of control (left) and PRRC2B knockdown cells (right) without DOX induction. (**F**) The quantification of relative cell number across different cell cycle stages in (E). (**G**) Representative flow cytometry images of control (shCtrl) and PRRC2B knockdown cells (shPRRC2B-328, shPRRC2B-626) 48 hours after DOX induction. (**H**) The quantification of relative cell number across different cell cycle stages in (G). For all the line plots and histograms, biological triplicates were performed. For all flow cytometry results, biological triplicates were performed, G0/G1, S, and G2/M peaks were autodetected and quantified by FlowJo, and percentages of total cells detected by flow cytometry were reported. Data were expressed as mean ± SD, and the Student’s *t* test was used to calculate the significance. NS, not significant; * *P* < 0.05; ** *P* < 0.01; *** *P* < 0.001.

**Figure S5.**
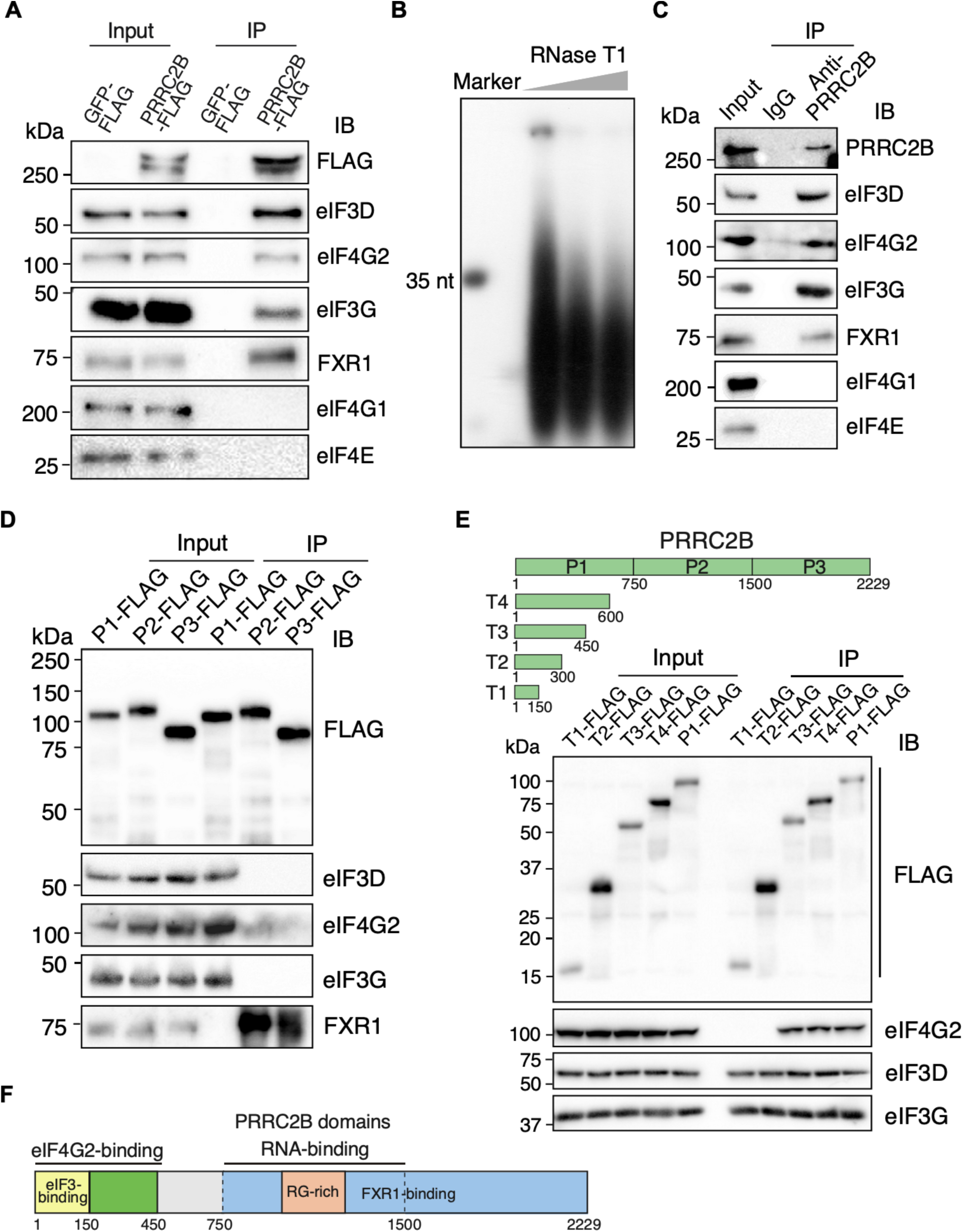
Co-immunoprecipitation of PRRC2B with translation initiation factors. (**A**) Western blot detection of the co-immunoprecipitation (Co-IP) of FLAG-tagged PRRC2B together with translation initiation factors (eIF4G2 and eIF3) and FXR1 without RNase T1 treatment. (**B**) SYBR gold staining shows the fragmentation of RNA in total cell lysates by 1 U/μl, 5 U/μl, and 10 U/μl RNase T1. (**C**) Western blot detection of the Co-IP of endogenous PRRC2B together with translation initiation factors and FXR1 upon 5 U/μl RNase T1 treatment. (**D**) Western blot detection of the Co-IP of FLAG-tagged PRRC2B fragments (P1, P2, P3) together with translation initiation factors and FXR1 upon 5 U/μl RNase T1 treatment. (**E**) Upper: Schematic of the fragmentation of the P1 region of PRRC2B protein. Truncated P1 (T1-T5) was made by removing a series of 150 amino acids from the C-terminus of P1. Lower: Western blot detection of the Co-IP of FLAG-tagged truncated P1 fragments (T1-T5) together with translation initiation factors upon 5U/μl RNase T1 treatment. (**F**) Schematic of the regions of PRRC2B protein interacting with eIF3, eIF4G2, FXR1, and RNA. Western blots were representative of biological replicates. Western blot results are representative of >2 independent experiments in (A, C, D, E).

**Figure S6.**
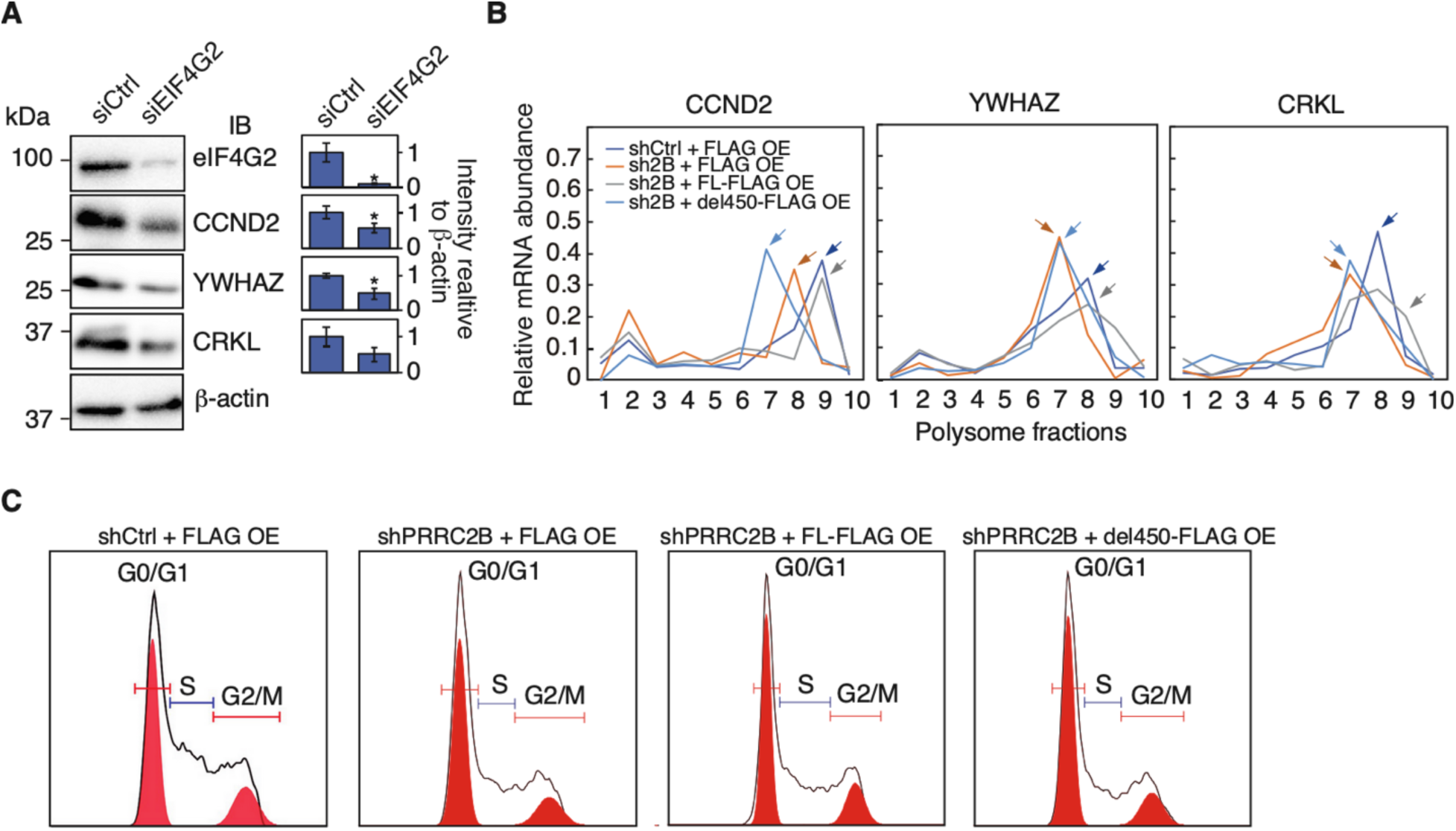
eIF4G2/eIF3-PRRC2B interaction is essential for translation of PRRC2B mRNA targets. (**A**) Western blot detection of the abundance changes of proteins (CCND2, YWHAZ, CRKL) in cells transfected with control siRNA with no target in human cells (siCtrl) and cells transfected with siRNA targeting CDS of *EIF4G2* mRNA (siEIF4G2). (**B**) mRNA abundance distribution (detected by RT-qPCR and represented as a percentage of total RNA) of PRRC2B-bound mRNAs (*CCND2*, *YWHAZ*, *CRKL*) in different polysome profiling fractions from cells under different conditions. Arrows indicate the peak of mRNA abundance. Biological duplicates were performed, and representative data were shown. (**C**) Representative flow cytometry images of cells under different conditions. For all flow cytometry results, biological triplicates were performed. β-actin is used as an internal control for western blot. For all assays involving PRRC2B knockdown, experiments were performed 36 hours after induction of PRRC2B knockdown. Western blots were representative of >2 biological replicates in (A). All quantitative western blot results are presented with quantification plots and analyzed by Student’s *t* test. Significance was calculated by comparing to siCtrl in (A). * *P* < 0.05; ** *P* < 0.01; *** *P* < 0.001.

**Extended Data 1.**
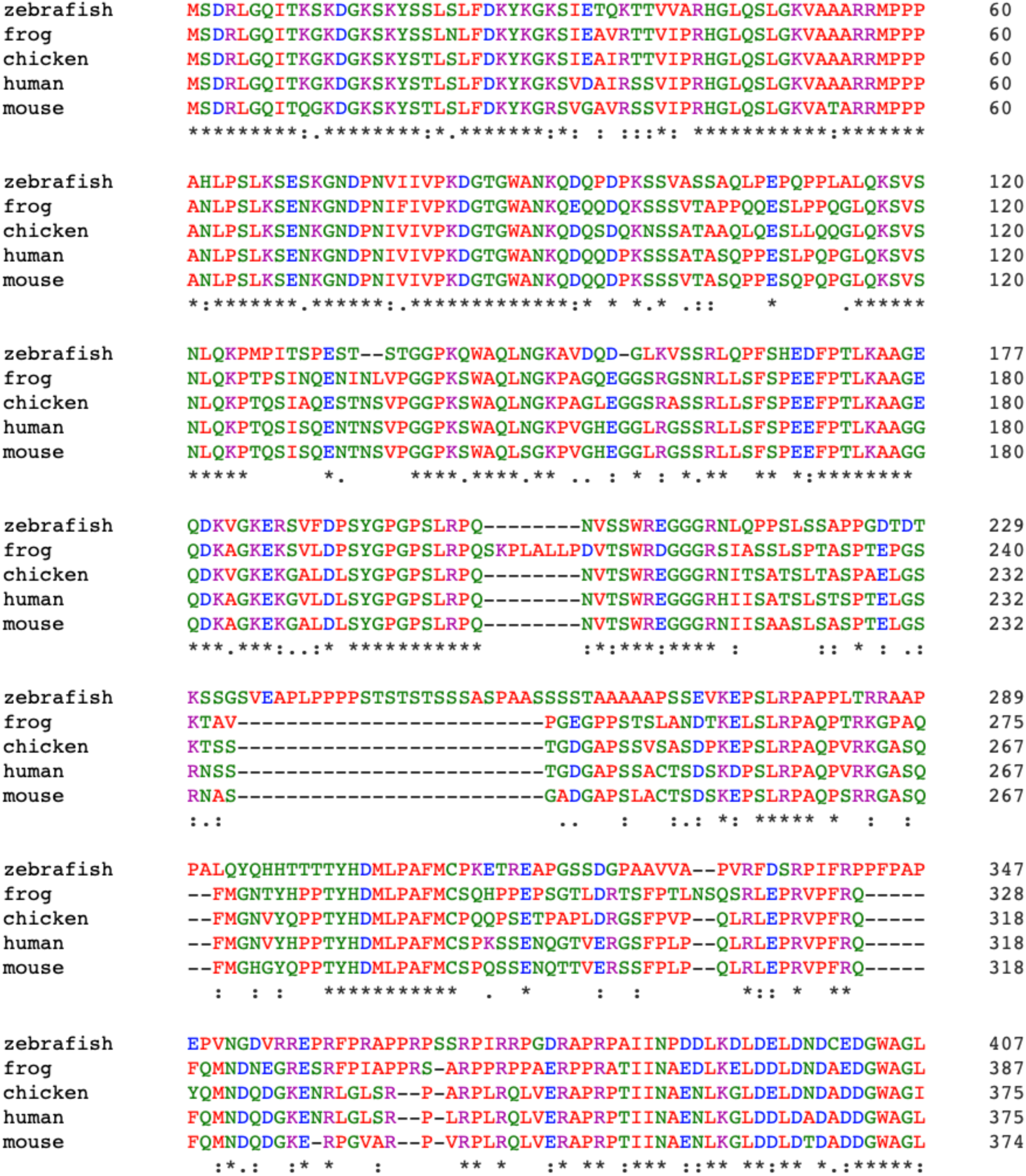
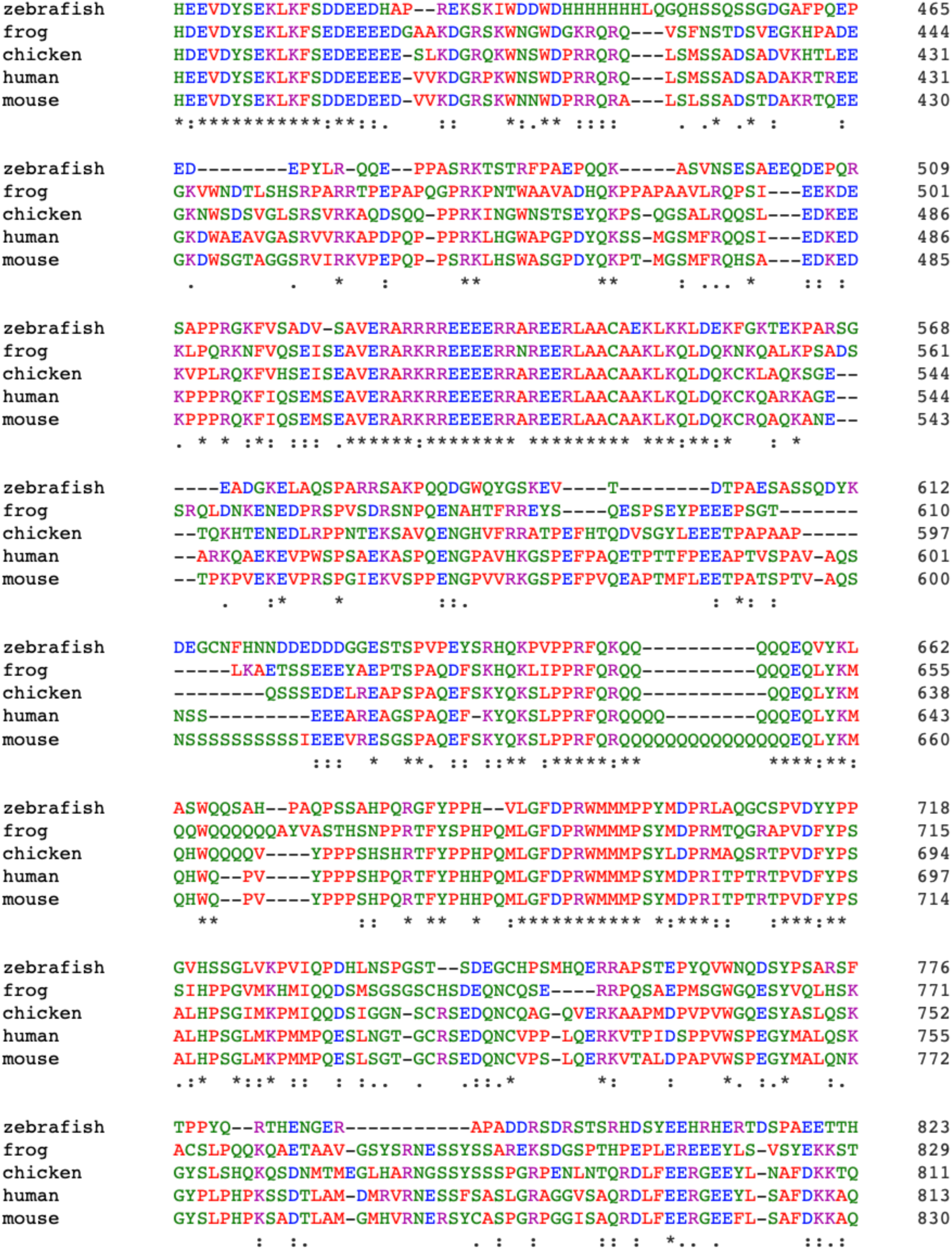
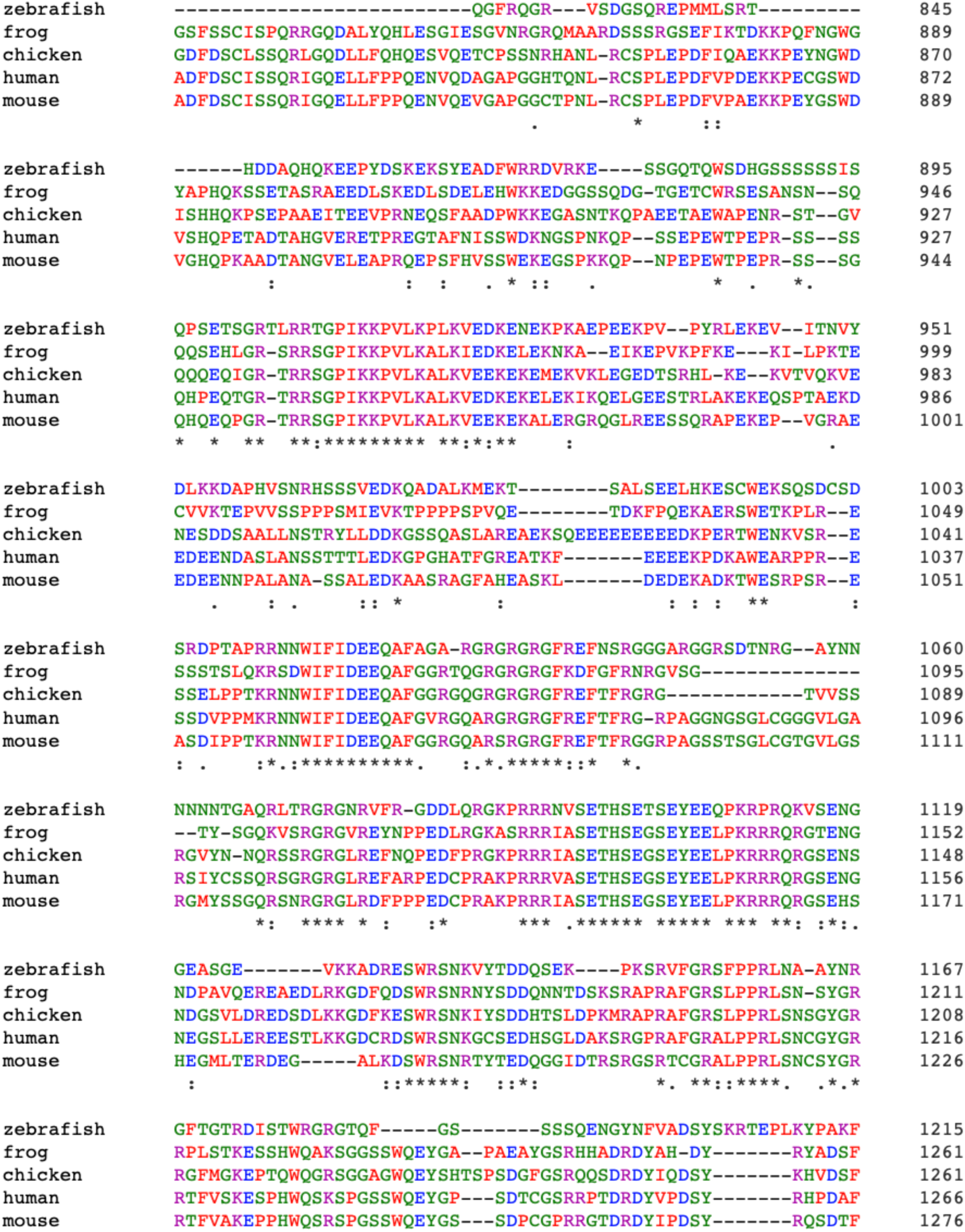
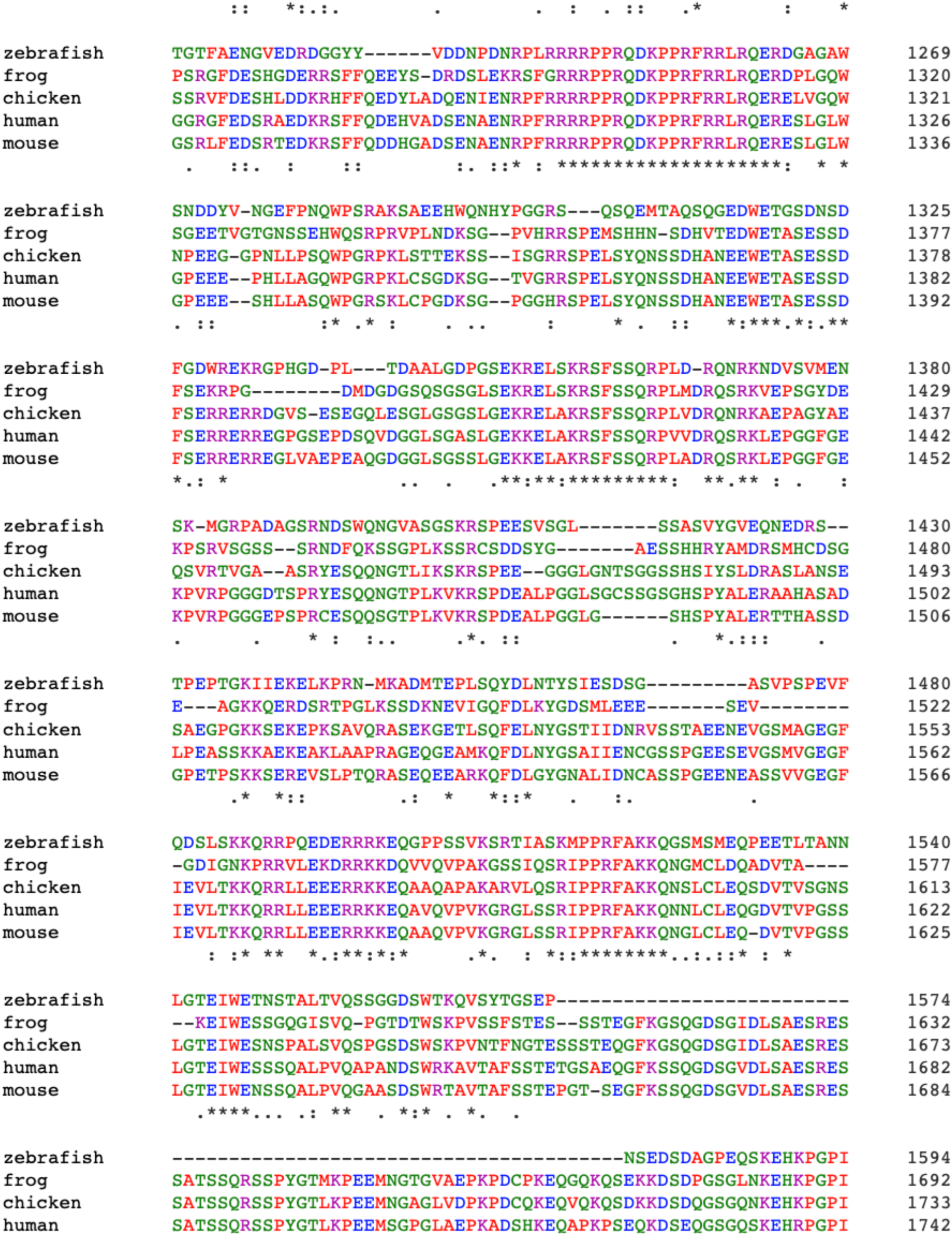
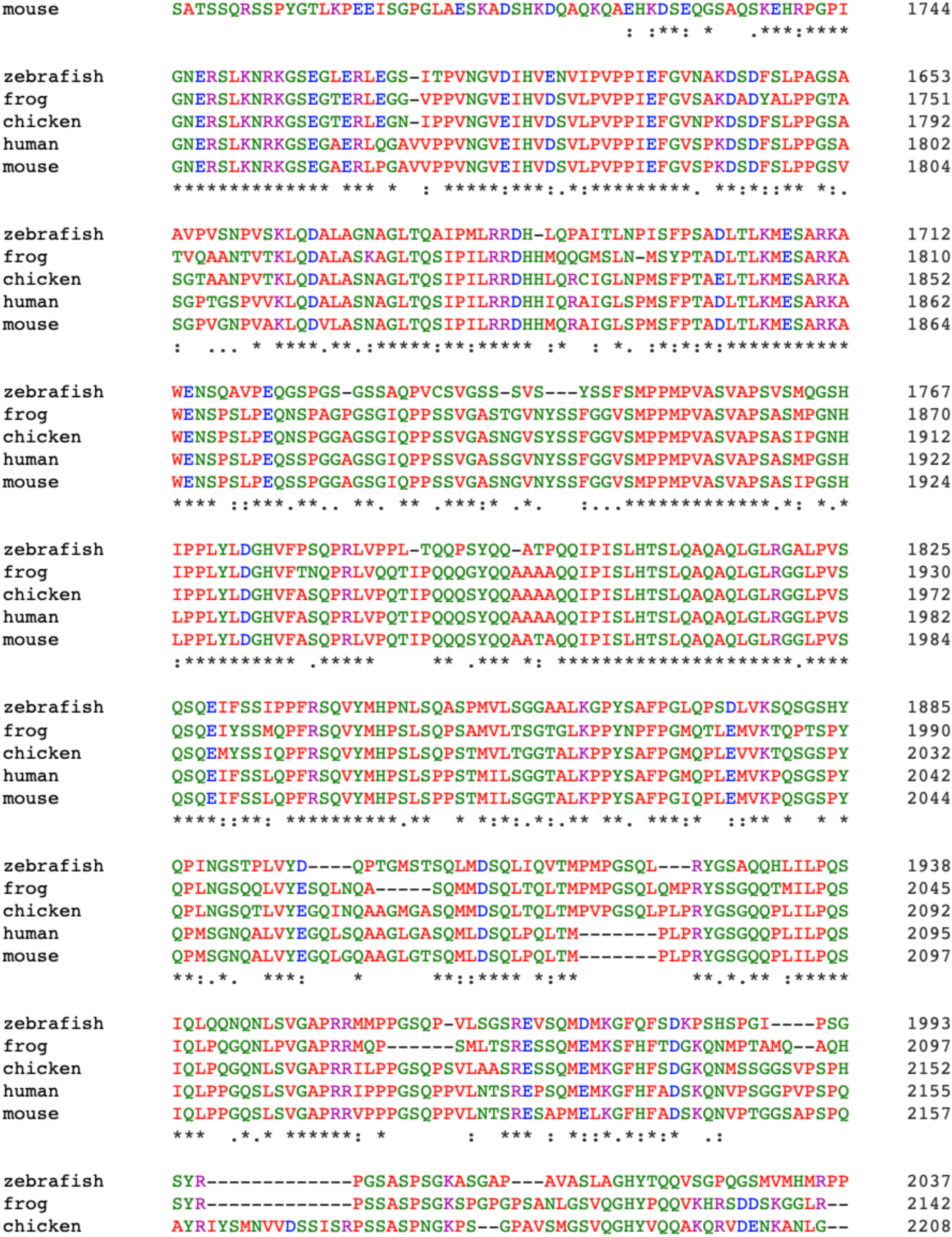
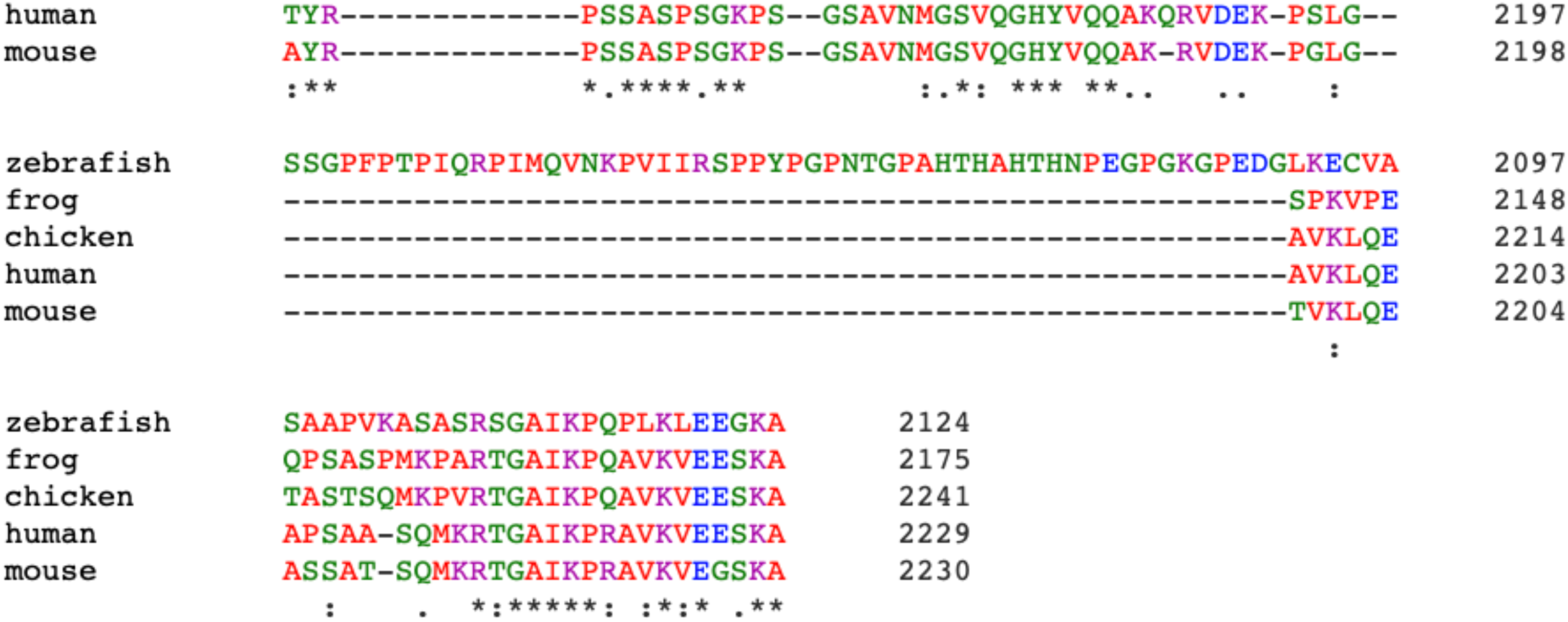
Sequence alignment of PRRC2B protein in five representative species was shown. PRRC2B gene is conserved in 460 species, including all kinds of mammals, reptiles, amphibians, and fish (e.g., zebrafish and shark) based on sequence information from NCBI.

**Extended Data 2.**
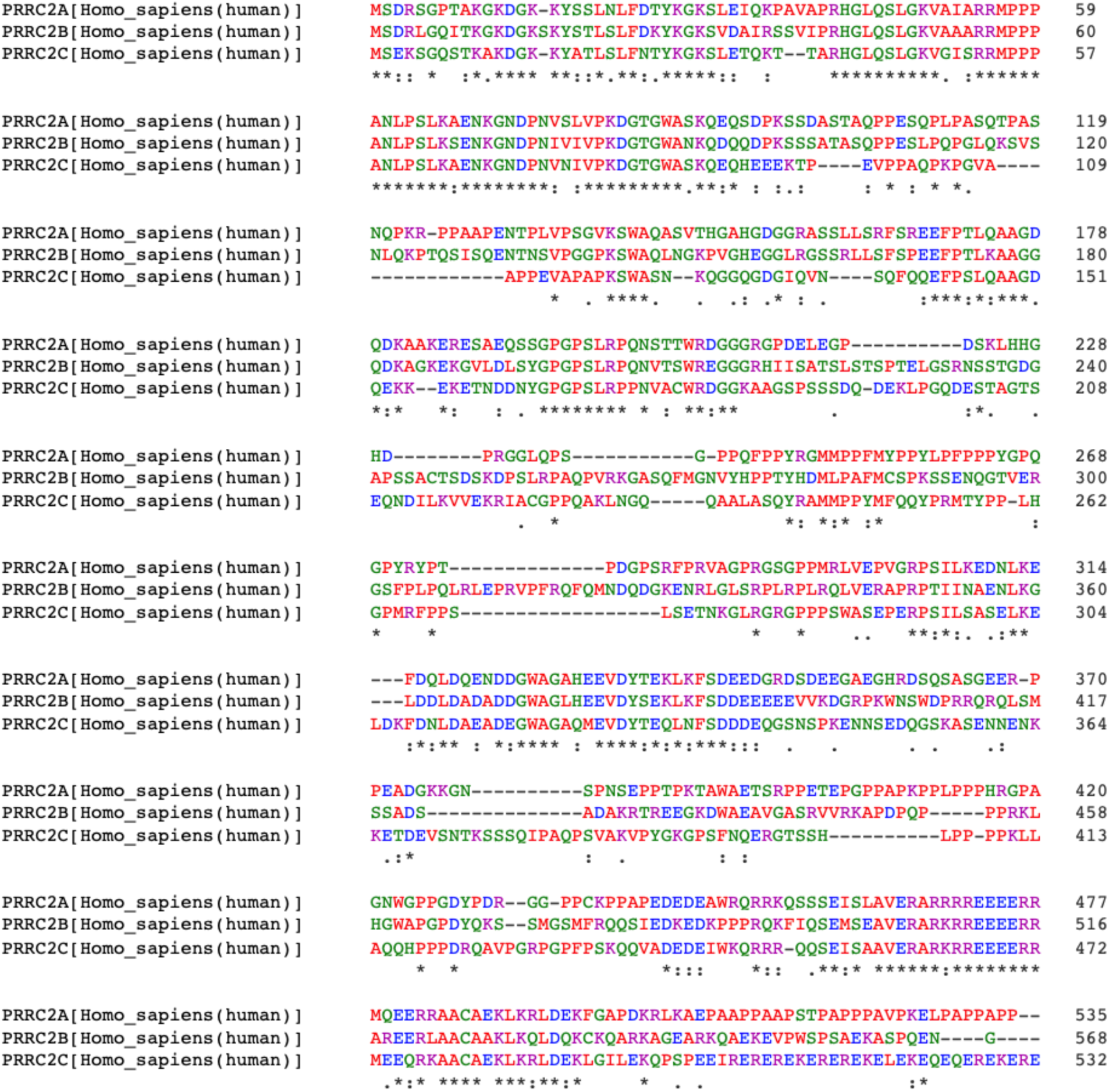
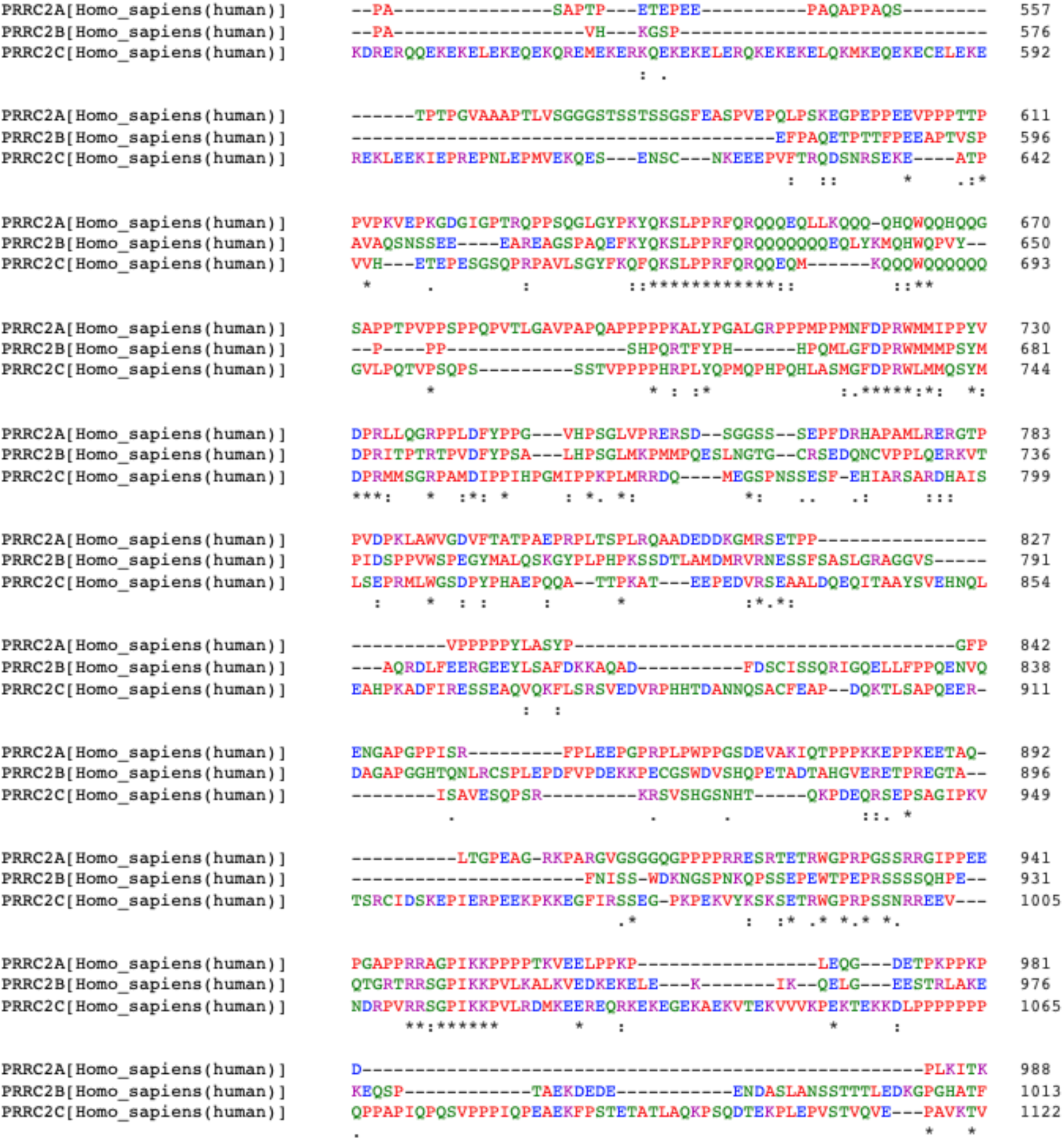
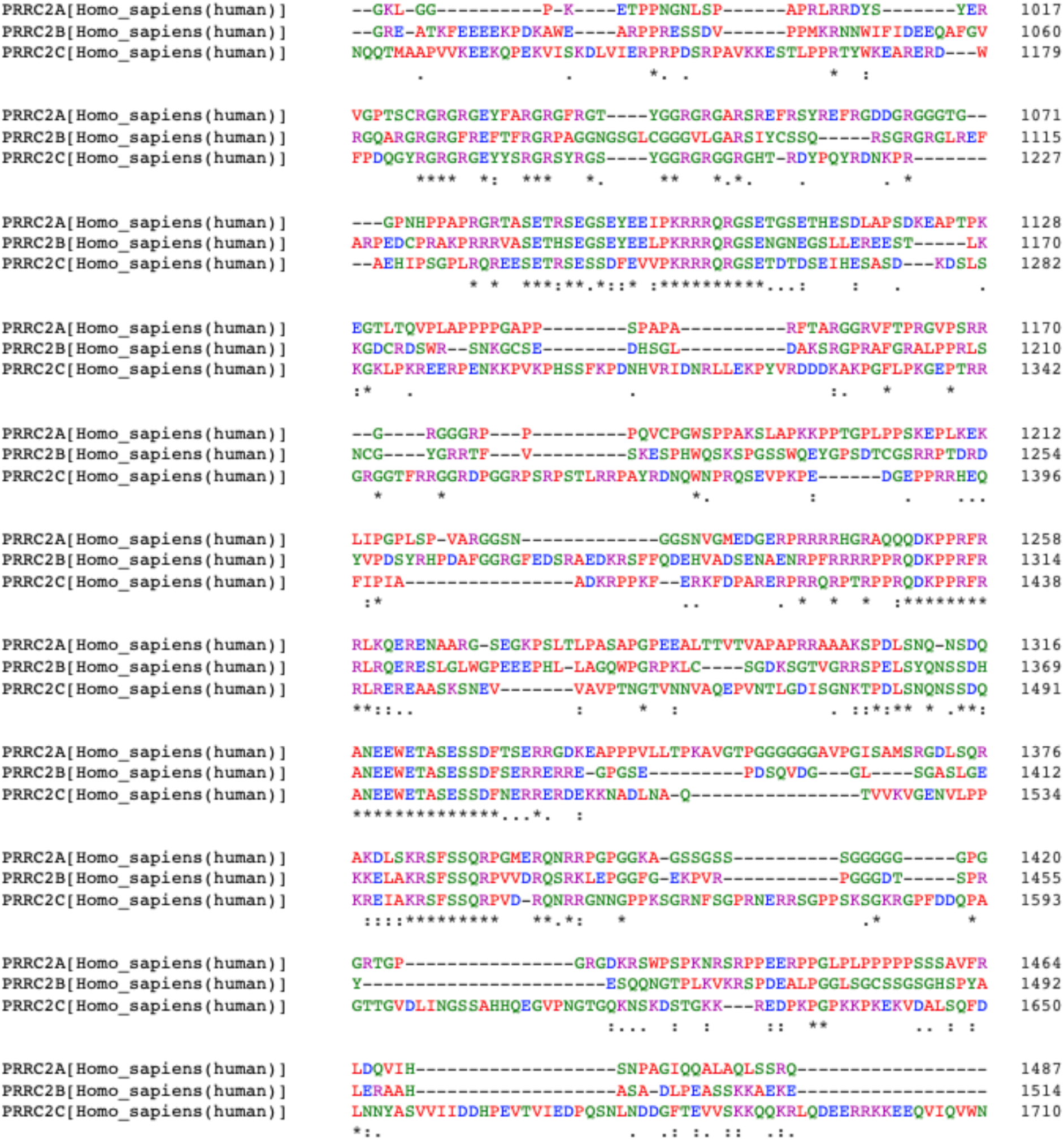
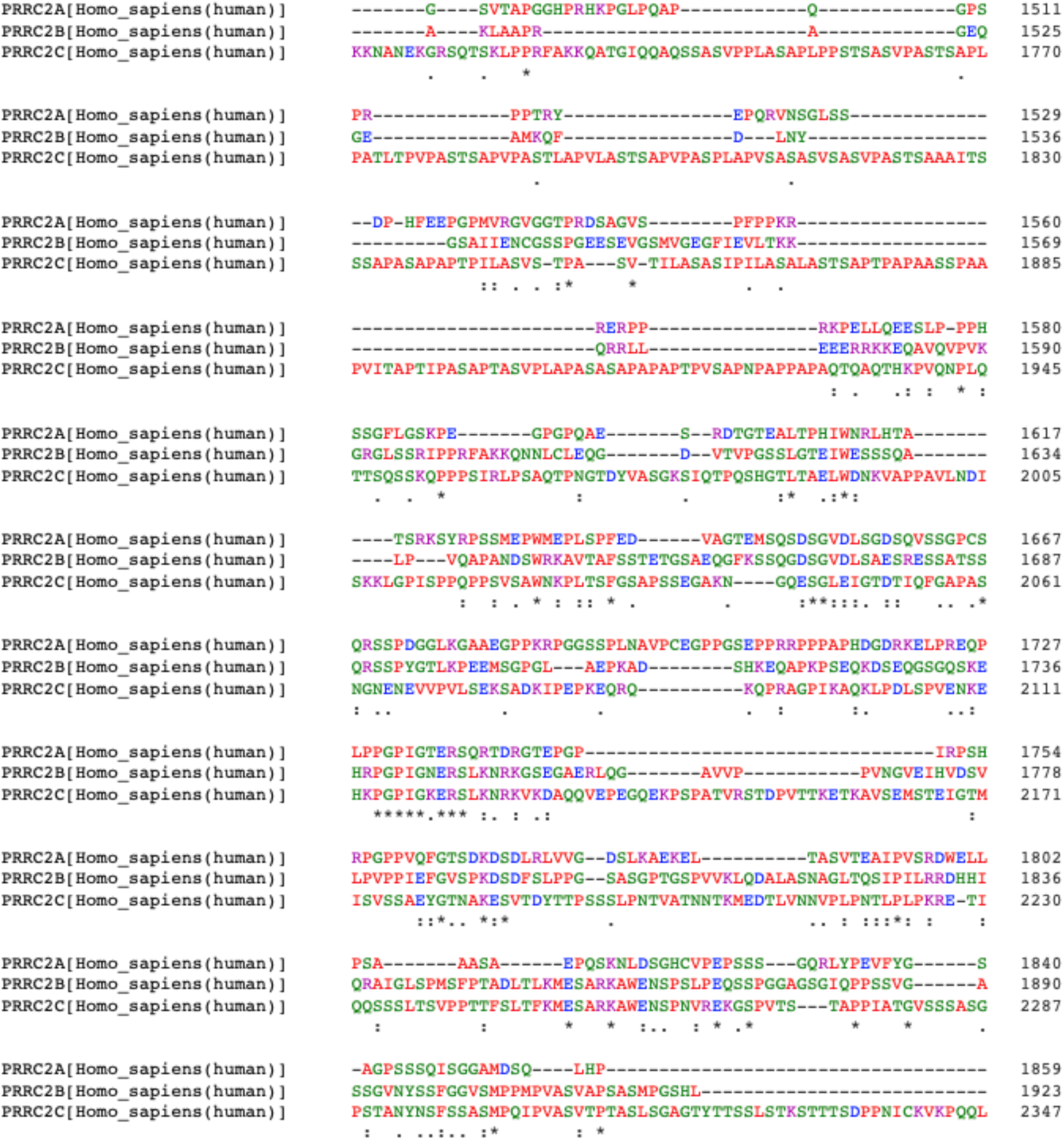
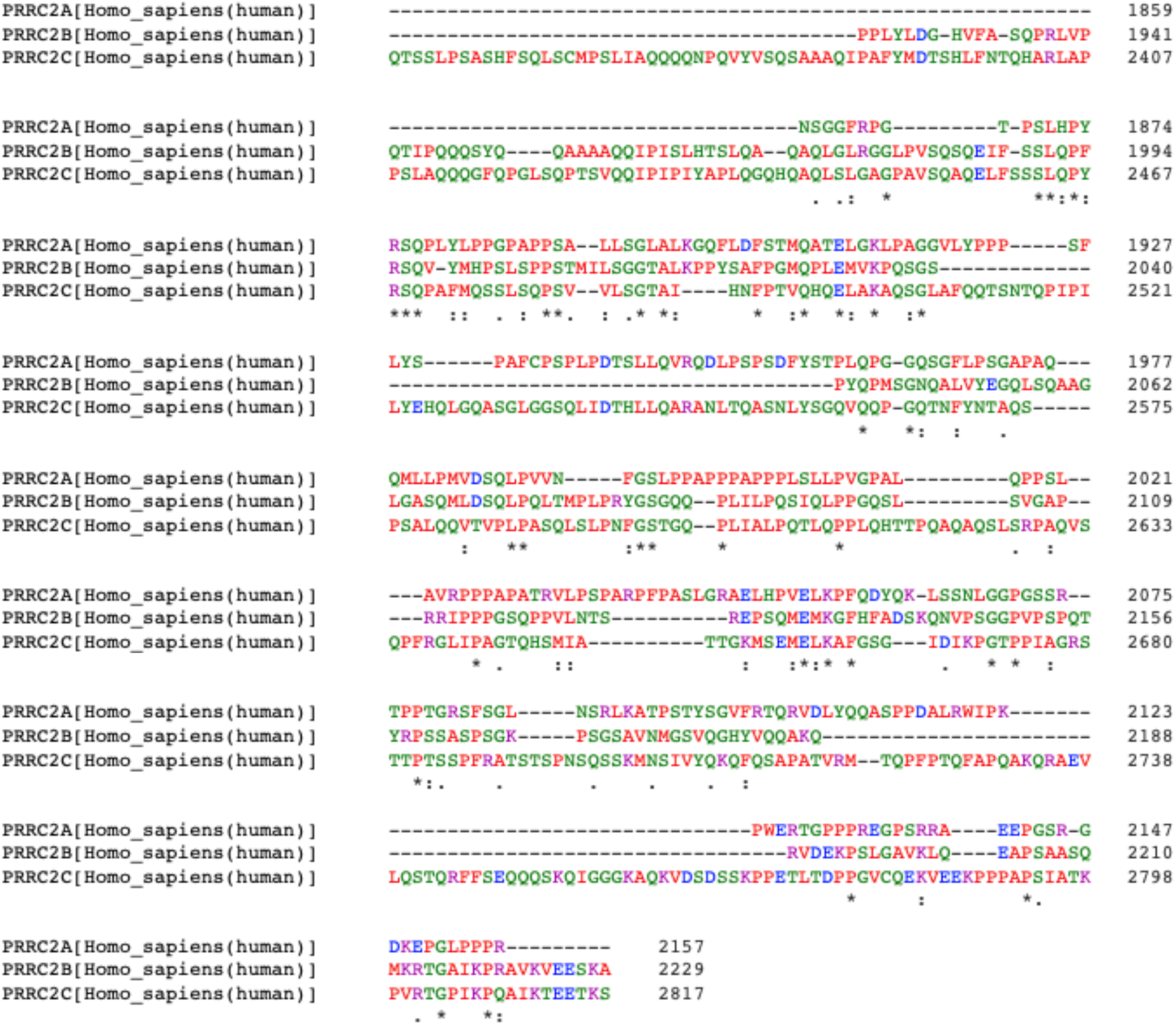
Sequence alignment of PRRC2A, PRRC2B, and PRRC2C proteins in humans was shown.

### Supplemental Material and Methods

#### Reagents, antibodies, plasmids, and siRNAs

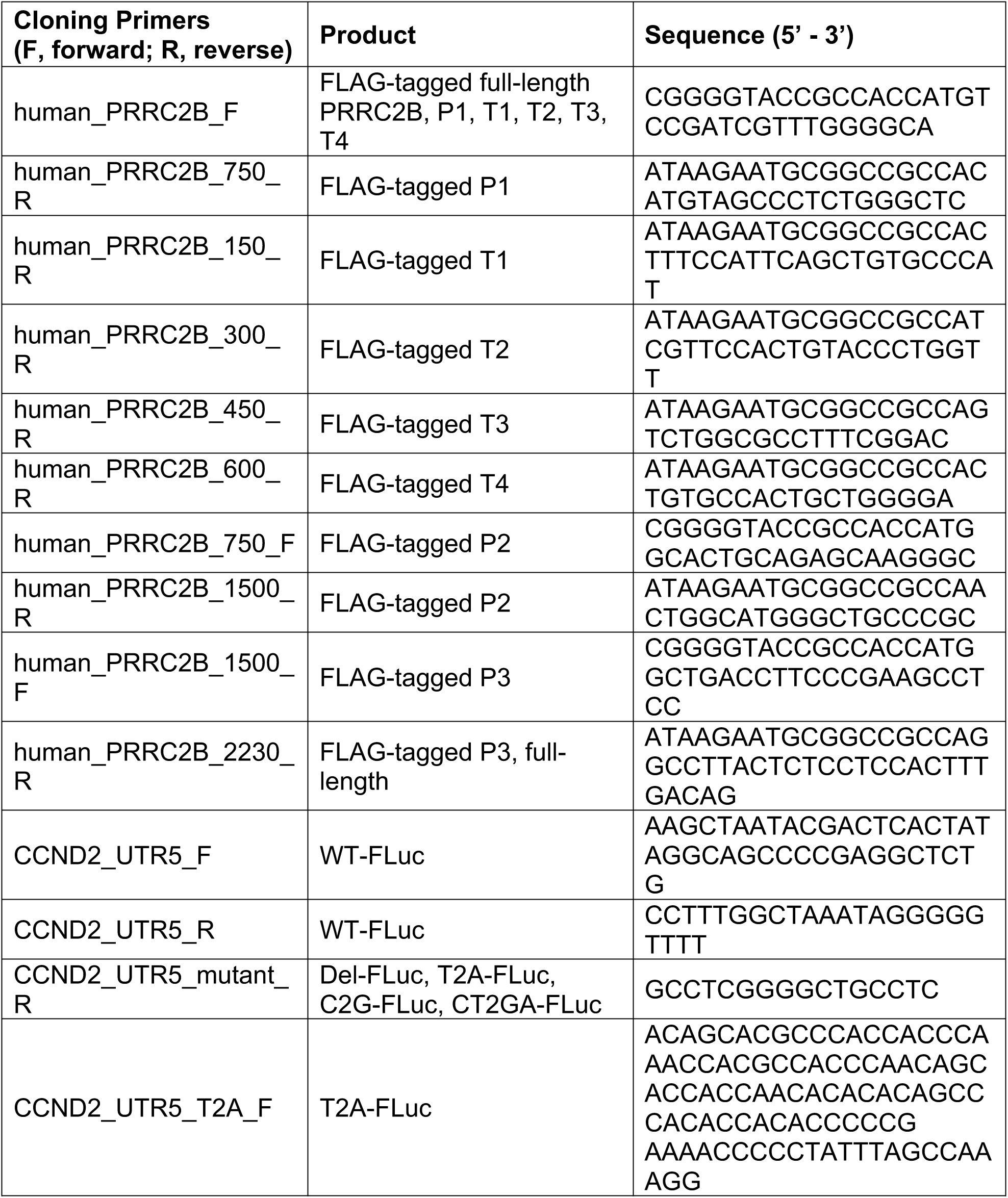
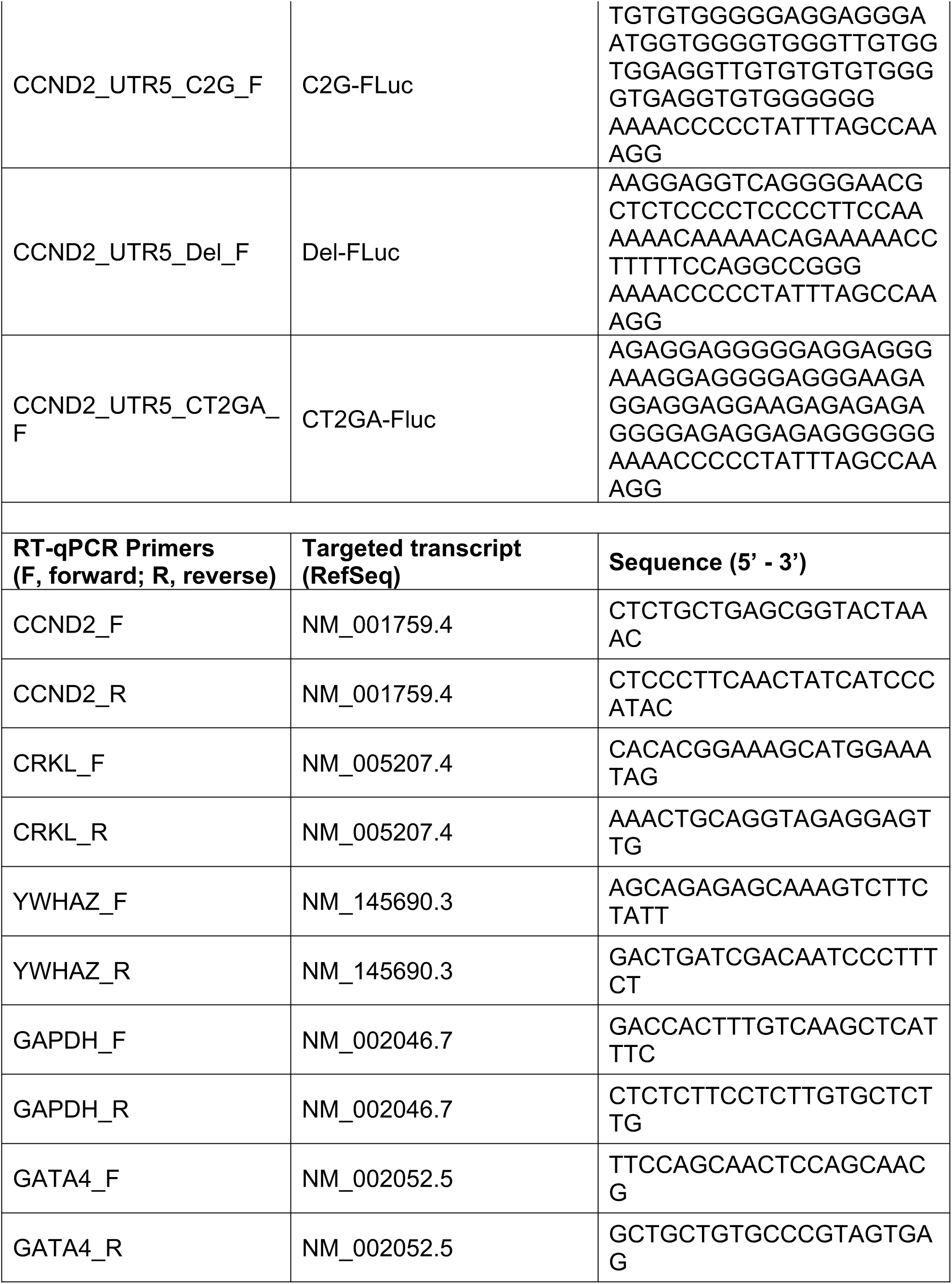
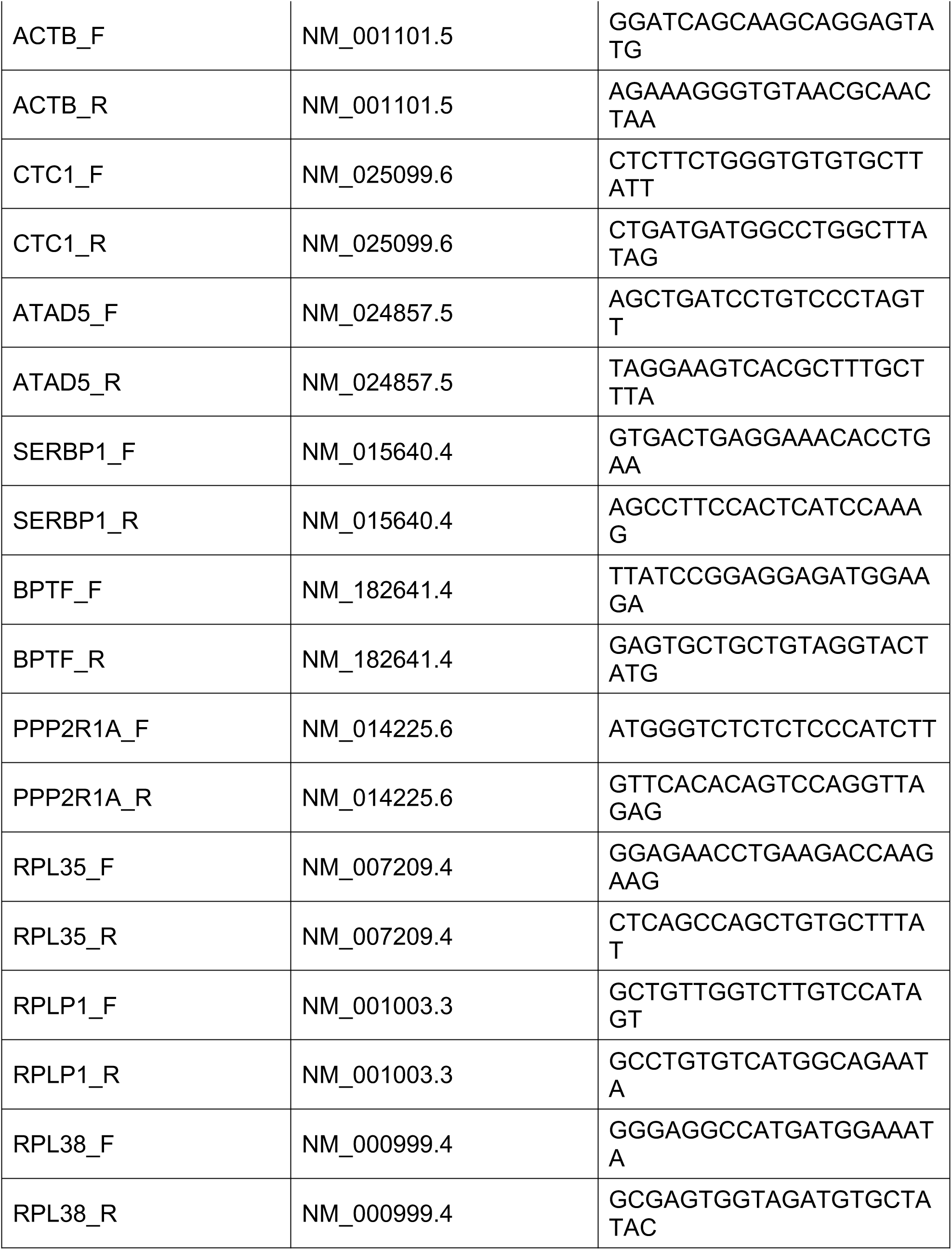
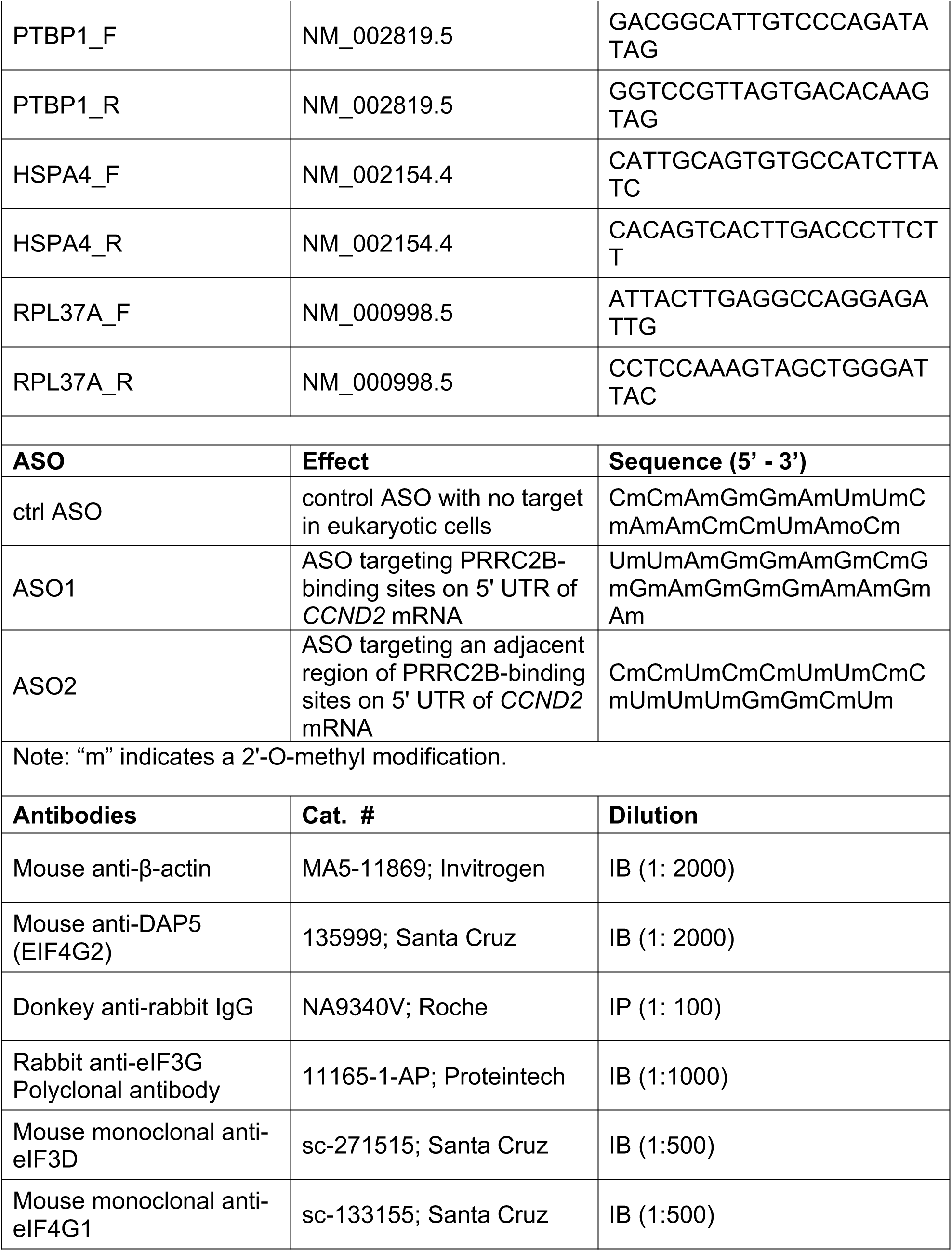
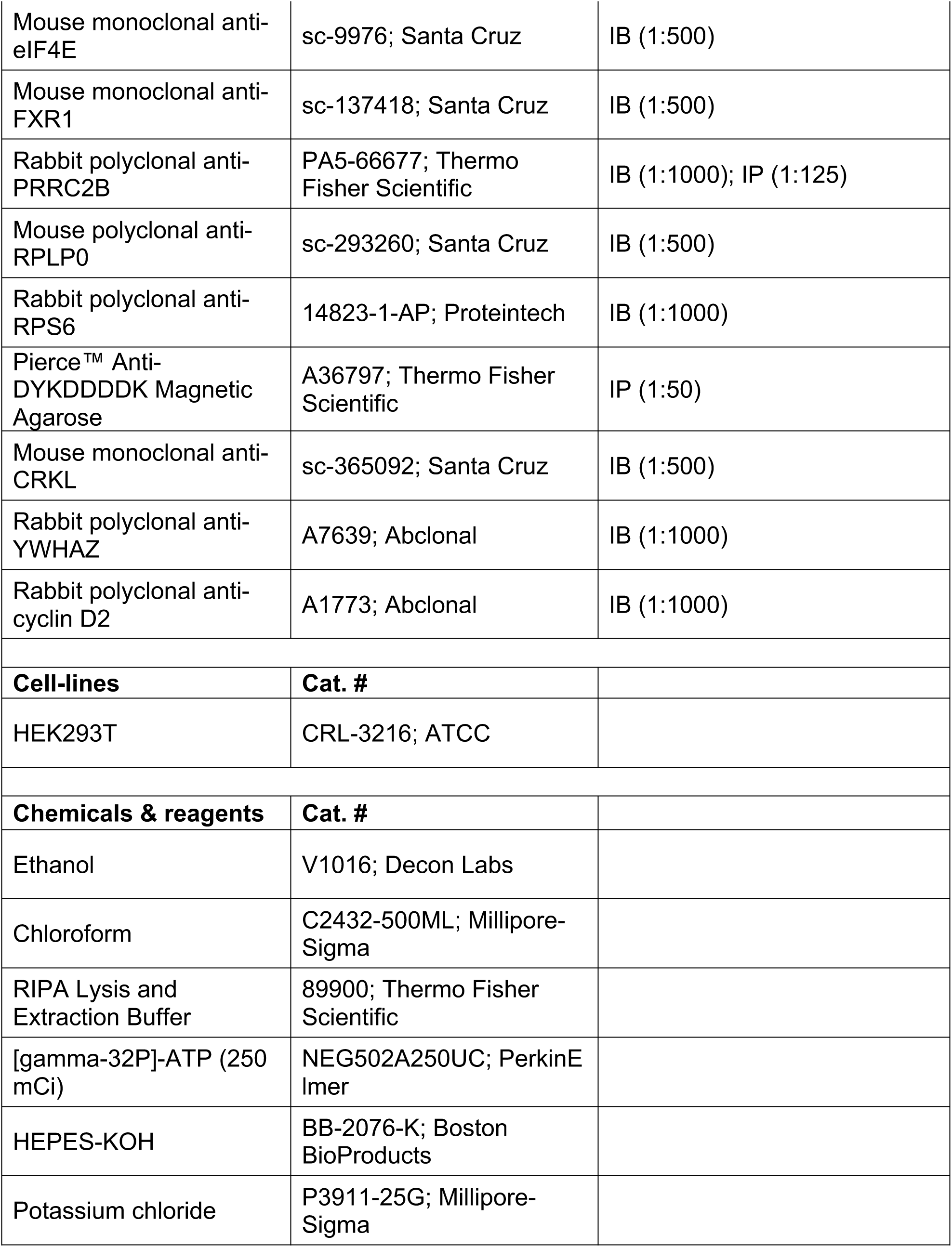
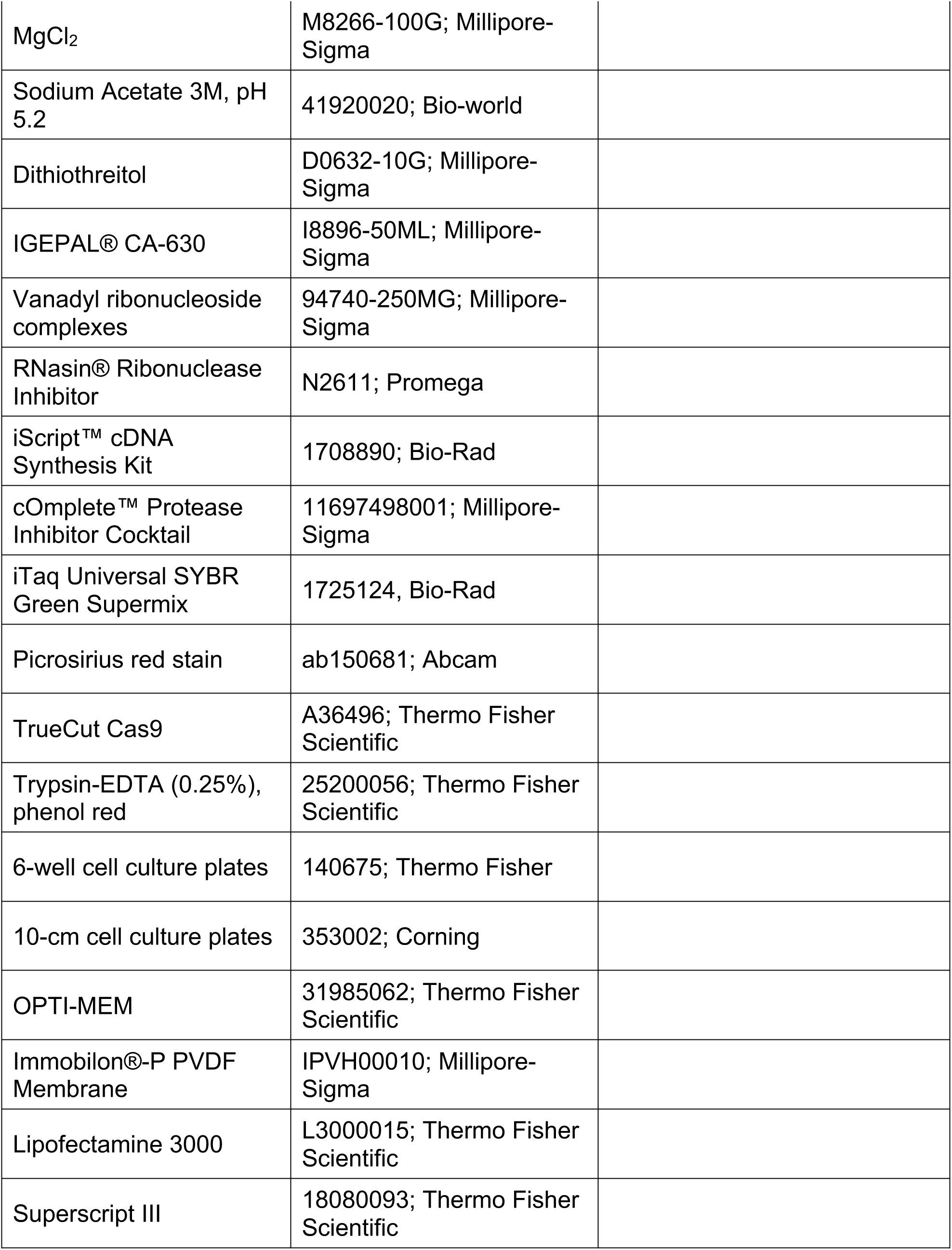
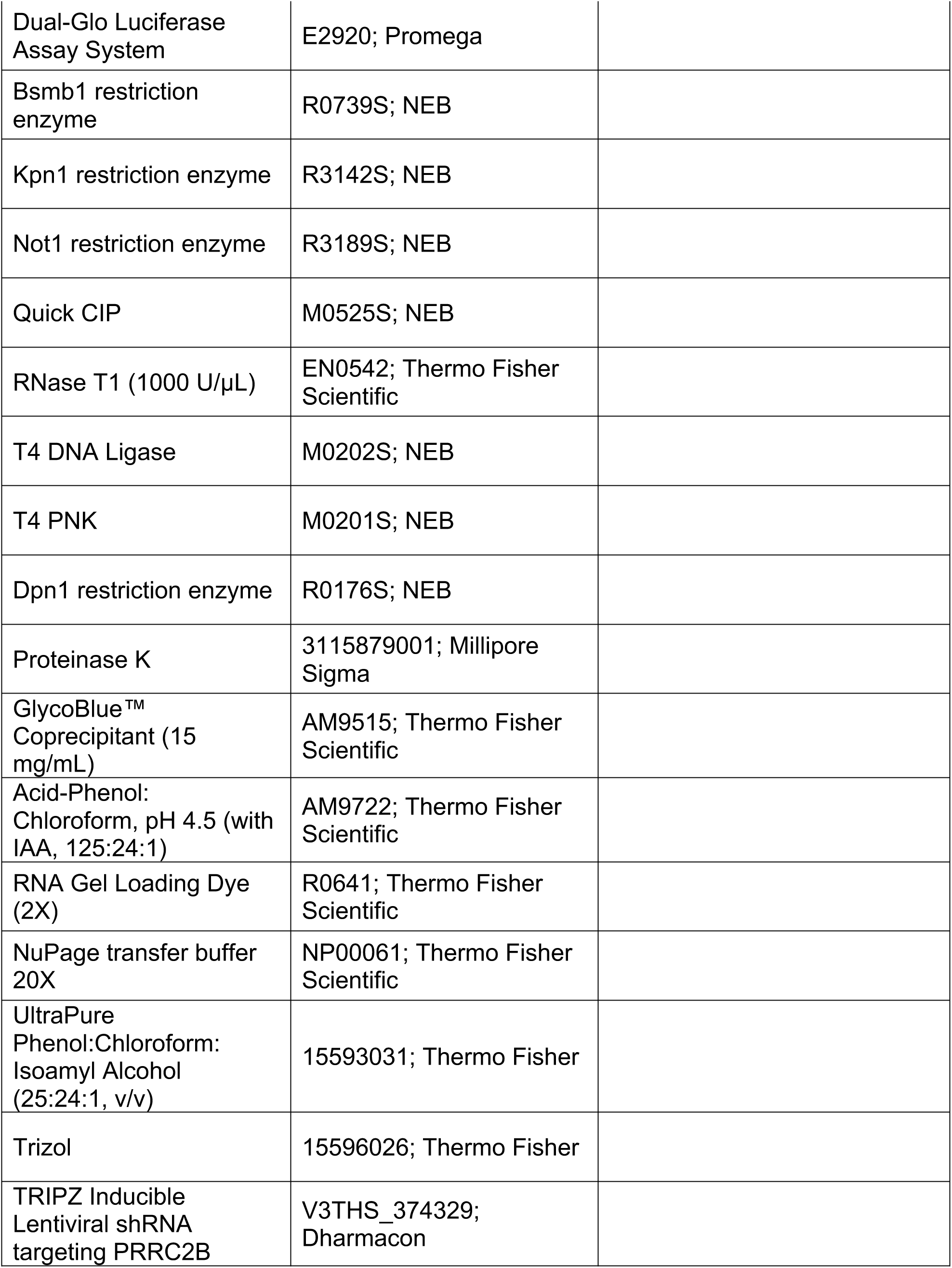
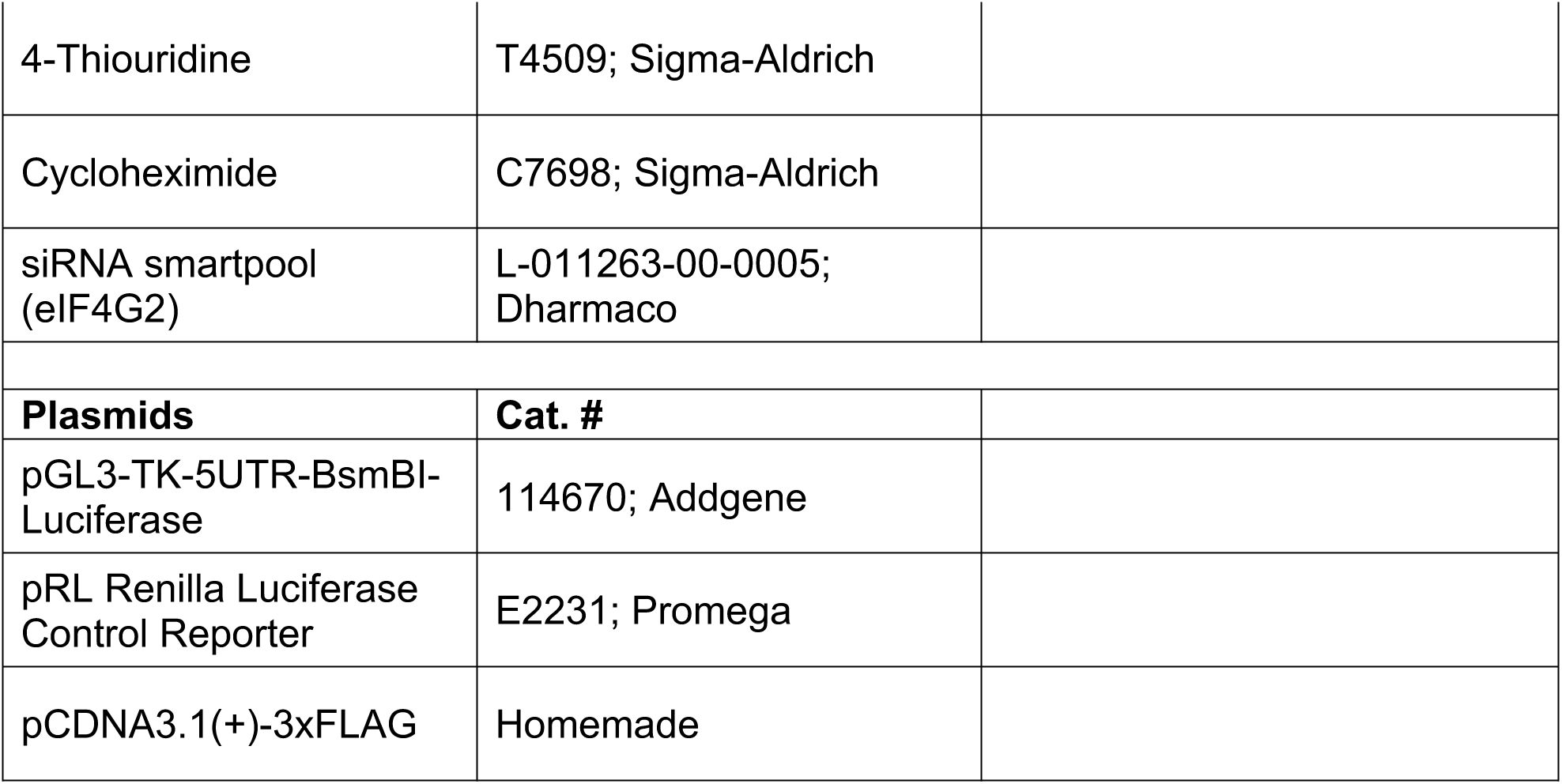

#### MIQE for RT-qPCR

For all RT-qPCR experiments in this study, > 1×10^9^ HET293T cells transfected with shRNAs, antisense oligos (ASOs), or luciferase reporters were used as experimental and control groups. All cell samples were obtained from cultured cells without dissection. If not used immediately, samples were snap-frozen in liquid nitrogen and stored at −80°C for up to 12 months.

For RNA extraction, cells were lysed with 1000 μl of Trizol (Qiagen) and mixed with 200 μl chloroform. The mixture was spun down at 16,000 g for 10 min. RNA was precipitated from the aqueous layer by adding two volumes of isopropanol and spinning down at 16,000 g for 10 min. The pellet was washed twice with 70% ethanol, left to dry, and resuspended in nuclease-free water. Potential genomic DNA contamination was removed by incubating with DNase I at 37°C for 10 min, followed by another round of RNA purification using chloroform: phenol: isoamyl alcohol, similar to those mentioned in the method section. Purified RNA was subject to 1% agarose gel electrophoresis to check for RNA integrity (reflected by 28S/18S > 1.0) and genomic DNA contamination. RNA quantification was performed by measuring A260 using Nanodrop (Thermofisher ND-ONE-W). RNA purity was assessed by A260/A280. Inhibition of reverse transcription was tested using 1000 ng, 500 ng, and 250 ng of RNA. Cq values of the inhibition test are shown in **Table S8**.

cDNAs were prepared using iScript master mix RT Kit (Bio-Rad) following the manufacturer’s suggestions. Briefly, 500-1000 ng of RNA were added to a 20 μl reaction containing reverse transcriptase, RNase inhibitor, dNTPs, primers (optimum blend of oligo(dT) and random primers), MgCl_2_, and stabilizers. Reactions were performed at the following conditions: (priming, 5 min at 25°C; reverse transcription, 20 min at 46°C; RT inactivation, 1 min at 95°C). No-RT control was included to ensure the synthesis of cDNA.

All qPCR primers were designed by the IDT qPCR primer design tool with no multiplex and synthesized as single-stranded DNAs by IDT with no extra modification. All primers target the exons in the coding region of the main open reading frame of the longest mRNA transcript (isoform) of each gene of interest. Detailed primer information and target sequence accession number are listed in Supplemental Material and Methods. All PCR amplicons are less than 200 nt long.

RT-qPCR-amplification was performed using SYBR Primer Assay kits (Bio-Rad 3 1752124) following the manufacturer’s suggestions. 10 μl reaction was set up in Bio-Rad Hard-Shell® 96-Well PCR Plates (#HSP9601) with 1 μl primer (10 μM), 5 μl Bio-Rad iTaq Universal SYBR Green Supermix (containing an optimal amount of Mg^2+^, hot-start iTaq DNA polymerase, and dNTP), 0.2 μl of cDNA from a 20 μl RT reaction, and RNase-free water. qPCR was performed using Bio-Rad CFX Connect Real-Time PCR Detection System with the following program: 1 min at 95°C, (10 s at 95°C, 40 s at 60°C, read plate) x 40 cycles, 10 s at 95°C, melt Curve 65 – 95°C, increment 0.5°C for 5 s, plate Read.

Cq values were determined by the regression mode in Bio-Rad CFX Connect Real-Time PCR Detection System. Cq values < 40 (no-template control has Cq value > 45) are considered reasonable. For each primer set, the identity of the resulting PCR product was confirmed by cloning and Sanger sequencing. Melting curves were used in each subsequent PCR to verify that each primer set reproducibly and specifically generates the same PCR product. A standard curve for each primer set was generated using 1x, 2x, 2.5x, 4x, 8x, 16x, 25x, 250x, 2500x, and 25000x diluted RNA (**Table S8**). Linear models were fitted using Microsoft Excel to calculate slope, y-intercept, and PCR efficiency in order to determine the Linear range of detection. At least biological duplicates and technical duplicates were performed for each measurement. Measurements with Cq variant (intra-assay) greater than 1 were repeated.

For data analysis, relative mRNA abundance was calculated by the Livak-Schmittgen method {2}^{-\Delta \Delta Cq}. Normalization was performed against 18S rRNA (internal control) based on its constant high expression in all samples or Renilla mRNA spike-in (if used). All quantitative data were presented as mean ± SD and analyzed using Excel (Microsoft Office). Statistical analyses were performed using the two-tailed Student’s *t* test with *P* < 0.05 considered significant.

## Notes

### Competing Interest Statement

The authors have declared no competing interest.

### Summary of Updates

This revised version was updated based on the reviewers comments from Nucleic Acids Research journal. Here are the major changes. The shRNA expressing vector we used has a red fluorescence protein (RFP) co-expressed with shRNA upon Dox treatment to indicate successful induction. We observed the RFP signal under a microscope after a 12-hour overnight incubation of doxycycline (Dox). In the original version of the manuscript, we considered this the start time point of a successful induction, and all the labels in the manuscript are actually counted after the 12-hour overnight incubation of Dox. Therefore, if we count from the time when Dox is initially added, the 6 hrs label becomes 18 hrs. Based on this fact, we corrected the timeline labeling to count from the time when Dox was initially added throughout the entire manuscript. This will affect Figure 2, S2, 3, S3, 4, S4, 6, S6. This will not affect the samples labeled 0 hrs in Figure 2H since no Dox is added at all for those samples. Given the promiscuity of shRNA, we have also rigorously repeated the experiments in Figure 2H and Figure 3 using two additional shRNA targeting different regions of PRRC2B main ORF. (shPRRC2B-626 and shPRRC2B-328). Detailed information about these shRNAs has been added to the method section. Although these two shRNAs (especially 328) do not knockdown PRRC2B as well as the one (shPRRC2B-329) used in Figure 2H, we were able to recapitulate the changes in translation, cell proliferation, and cell cycle progression (at a later time point due to lower knockdown efficiency) effects as shown in our updated Figure S2, S4. We added the quantification and statistics analysis data of western blots shown in Figures 2C, 2H, 3D, 6A, and 6D. We also included the quantification and statistical methods in figure legends and method sections of the main text.

